# Specialization of the Human Hippocampal Long Axis Revisited

**DOI:** 10.1101/2023.12.19.572264

**Authors:** Peter A. Angeli, Lauren M. DiNicola, Noam Saadon-Grosman, Mark C. Eldaief, Randy L. Buckner

## Abstract

The hippocampus possesses anatomical differences along its long axis. Here the functional specialization of the human hippocampal long axis was explored using network-anchored precision functional MRI (N = 11) paired with behavioral analyses (N=266). Functional connectivity analyses demonstrated that the anterior hippocampus was preferentially correlated with a cerebral network associated with remembering, while the posterior hippocampus was correlated with a distinct network associated with behavioral salience. Seed regions placed within the hippocampus recapitulated the distinct cerebral networks. Functional characterization using task data within the same intensively sampled individuals discovered a functional double dissociation between the anterior and posterior hippocampal regions. The anterior hippocampal region was sensitive to remembering and imagining the future, specifically tracking the process of scene construction, while the posterior hippocampal region displayed transient responses to targets in an oddball detection task and to transitions between task blocks. These findings suggest specialization along the long axis of the hippocampus with differential responses reflecting the functional properties of the partner cerebral networks.

## Introduction

Multiple lines of evidence suggest different possibilities for how the long axis of the hippocampus might be functionally specialized (Poppenk et al. 2013; Strange et al. 2014). As one example, anchoring from the rich literature on place fields, the dorsal / posterior hippocampus has been implicated in fine-scale spatial representation and navigation (O’Keefe and Dostrovsky 1971; O’Keefe 1976; Maguire et al. 1997, 1998, 2000; Ryan et al. 2010; Woollett and Maguire 2011, 2012) while the ventral / anterior hippocampus is suggested to support coarser spatial representations (Jung et al. 1994; Kjelstrup et al. 2008; Hirshhorn et al. 2012; Brunec et al. 2018). As another example, building from the association between hippocampal damage and declarative memory deficits (Scoville and Milner 1957; Milner et al. 1968; Zola-Morgan et al. 1986; Cohen and Eichenbaum 1993), the anterior hippocampus has been reported to be preferentially involved in memory encoding while the posterior hippocampus is involved in retrieval (Lepage et al. 1998; Poppenk and Moscovitch 2011). However, other studies have found minimal effect in the posterior hippocampus (Spaniol et al. 2009) or even the opposite specialization (Schacter and Wagner 1999). These investigations and others support differential specialization along the long axis, but do not agree on a framework to explain the nature of the specialization.

Building on the above work, one approach to understanding long axis specialization examines how hippocampal subregions connect to distinct cerebral networks (e.g., Ranganath and Ritchey 2012). Anatomical tracing studies in non-human primates note differential patterns of extrinsic connectivity along the hippocampal long axis (Amaral and Witter 1989; Insausti and Amaral 2008; Witter and Amaral 2021), suggesting that the anterior and posterior hippocampus may be connected to different cerebral networks via the entorhinal cortex. These network connectivity findings provide context to motivate functional explorations. Specifically, if subregions of the hippocampus are linked to distinct cerebral networks, then functional specializations within the hippocampus might be clarified by exploring response properties related to the specializations of the partner cerebral networks.

Hippocampal-cortical connectivity can be estimated in humans using functional MRI (fMRI) connectivity (Biswal et al. 1995; Fox and Raichle 2007). Functional connectivity from the hippocampus and adjacent parahippocampal cortex reliably includes the retrosplenial / posterior cingulate cortex, caudal posterior parietal cortex, ventromedial prefrontal cortex, dorsolateral prefrontal cortex, and the rostral temporal cortex extending to the pole (e.g., Greicius et al. 2004; Vincent et al. 2006; Kahn et al. 2008; Frank et al. 2019; Barnett et al. 2021). The group-averaged human estimate of hippocampal-cortical network organization is similar to the distributed anatomical projection patterns in non-human primates (e.g., Buckner et al. 2008; Binder et al. 2009; Margulies et al. 2009; Buckner and Margulies 2019; see also Blatt et al. 2003), reinforcing the potential of a network-anchored approach to functional characterization (despite limitations; see Buckner et al. 2013; Murphy et al. 2013; Power et al. 2014 for discussion).

In a major advance using within-individual precision neuroimaging, Zheng et al. (2021) found that two distinct cerebral networks possess differential coupling to the anterior and posterior hippocampus. The first network, correlated with the anterior hippocampus, is topographically similar to canonical group-based estimates of the hippocampal-cortical network mentioned above and has been functionally associated with autobiographical remembering and scene construction (e.g., Svoboda et al. 2006; Hassabis and Maguire 2007; Schacter et al. 2007; Andrews-Hanna et al. 2014; DiNicola et al. 2020, 2023a, 2023b; Du et al. 2023). This network has also been identified within individuals using high-field fMRI from seed regions placed in the subiculum (Braga et al. 2019) and parahippocampal area TH (Reznik et al. 2023). By contrast, the posterior hippocampus is correlated with a distinct cerebral network that Zheng and colleagues (2021) described as the Parietal Memory Network (PMN; see Gilmore et al. 2015). This second network includes posterior midline regions that are spatially distinct from the anterior hippocampal network.

Adding nuance to inferring functions of the hippocampal subregions, the network coupled to the posterior hippocampus includes regions resembling another network extensively studied in the literature, generally referred to as the Salience Network (SAL; Seeley et al. 2007; Seeley 2019). By our estimation, the SAL and PMN networks may be the same network historically described by two different literatures (referred to as SAL / PMN; see Du et al. 2023). The observation that the posterior hippocampus may be coupled to a network which responds to transient events raises the possibility that the posterior hippocampus itself may support the detection of salience or novelty (e.g. Rolls et al. 1989; Knight 1996; Fyhn et al. 2002; Kumaran and Maguire 2009). That is, while the field has more often focused on examining hippocampal specialization in relation to spatial and mnemonic processing, the posterior hippocampus’ potential linkage to the SAL network suggests a counter-intuitive hypothesis: the posterior hippocampus may dissociate from the anterior hippocampus by its response to salient transient events (for background see Konishi et al. 2001; Fox et al. 2005; Dosenbach et al. 2006; Seeley et al. 2007; Seeley 2019).

With these possibilities in mind, the present work revisits the topic of hippocampal long axis specialization.

## Methods

### Overview

The goal of the present work was to explore functional specialization along the long axis of the hippocampus within individuals. Participants whose cerebral networks were previously estimated (Du et al. 2023) were reanalyzed here with focus on the hippocampus. Seeking to replicate Zheng et al. (2021), extensive resting-state fixation data were first used to estimate differential correlation to cerebral networks along the hippocampal long axis. Then, after identifying anterior and posterior hippocampal regions, functional response properties were examined during a task involving remembering and imagining future scenarios (collectively referred to as “Episodic Projection”) as well as tasks targeting low-level oddball detection and transitions between task blocks. All estimates were extracted within the anatomy of individual participants and then averaged afterwards to avoid anatomical blurring. A key additional feature of our approach, building from DiNicola et al. (2023a), was to obtain behavioral assessments of the task strategies employed during the Episodic Projection task trials to further inform the nature of the component processes that drive the hippocampal response.

### Participants

English-speaking adults (ages 18-34) without neurological or psychiatric illness completed MRI scanning sessions (data from Du et al. 2023). After exclusions to ensure data quality, the MRI data set included 11 participants with a mean age of 22.4 yr (SD = 4.1; 11 right-handed; 8 women). Participants were from diverse racial and ethnic backgrounds (7 of 11 individuals self-reported as non-white and/or Hispanic). Behavioral data used to estimate task trial components included, after exclusions for quality control, 266 English-speaking adults (ages 20-28), located within the United States with high ratings for completion and performance (90%+ approval rating with at least 100 prior tasks approved), who contributed data online through Amazon Mechanical Turk using Cloud Research (Litman et al. 2017). Behavioral participants had a mean age of 23.2 yr (SD = 1.9, 136 women). All participants were paid and provided informed consent according to a protocol approved by the Harvard University IRB.

### Data Acquisition, Quality Control, and Preprocessing of MRI Data

Scanning was conducted at the Harvard Center for Brain Science using a 3T Siemens MAGNETOM Prisma^fit^ MRI scanner as described in Du et al. (2023). For P2, a 64-channel phase-arrayed head-neck coil (Siemens Healthcare, Erlangen, Germany) was used for 2 sessions, and a 32-channel phase-arrayed head coil (Siemens Healthcare, Erlangen, Germany) was used for the remaining 6 sessions. Data for all other participants used the 32-channel head coil, and the 64- and 32-channel coils were treated as comparable.

Participants completed 8-10 neuroimaging sessions with scanning generally completed over 6 to 10 weeks. Each session included multiple fMRI runs using a sequence sensitive to blood oxygenation level-dependent (BOLD) contrast (Kwong et al. 1992; Ogawa et al. 1992). Seventeen to 22 resting-state fixation runs (each 7 min 2 sec long) were acquired for each individual, to be used for functional connectivity analyses, along with sequence-matched task runs (see *Task Paradigms* below) and high resolution T_1_- weighted structural images (see Du et al. 2023 for additional information).

MRI data were examined for quality prior to analysis. Run-level exclusion criteria included: (1) maximum absolute motion exceeding 1.8mm, and (2) closed eyes during skipped or incorrect task trials, or poor task performance. Because each run of the Episodic Projection task was significantly longer than other tasks, the maximum absolute motion threshold for this task was raised to 2.5mm. Of the 15 recruited individuals, three participants (P5, P10, and P11) were excluded from analyses due to a high number of excluded Episodic Projection task runs or missed trials within the Episodic Projection task. One additional participant (P12) was excluded after preprocessing due to misregistration to the cortical surface. Usable resting-state fixation runs for included participants ranged from 16 (P2) to 22 (P7 and P13). Included task runs per participant (after additional behavioral exclusions as described in *MRI Task Paradigms*) ranged from 8 to 10 for the Episodic Projection task (5 total exclusions across participants), 3 to 5 for the Visual Oddball Detection task (3 total exclusions), and 8 for the Blocked Visual-Motor task (no exclusions). All data exclusions were finalized prior to analyses to prevent bias.

Preprocessing employed a custom analysis pipeline that aligned data across runs and sessions while minimizing spatial blurring (“iProc”, see Braga et al. 2019). Each participant’s data were registered to their own 1mm-isotropic native-space T_1_-weighted template and the MNI ICBM 152 1mm atlas, each through a single interpolation. Fixation data used for functional connectivity analysis had multiple nuisance variables regressed (6 head motion parameters as well as whole-brain, ventricular, and white matter signals and their temporal derivatives; 3dTproject, AFNI; Cox 1996, 2012), and underwent bandpass filtering at 0.01-0.10 Hz (3dBandpass, AFNI; Cox 1996, 2012). Task-based data had only whole-brain signal regressed. Resting-state fixation and task-based data were visualized in MNI space for hippocampal analyses (without smoothing). For all other analyses, data were projected from the T_1_-weighted native space to the fsaverage6 standardized cortical mesh using trilinear interpolation and a 2mm Full Width Half Maximum (FWHM) Gaussian smooth (40,962 vertices per hemisphere, see Fischl et al. 1999).

### Cerebral Networks Estimated on the Surface Within Individuals

Cerebral networks were identified based on functional connectivity using a multi-session hierarchical Bayesian model (MS-HBM) approach computed on the cerebral surface (Kong et al. 2019) (see Du et al. 2023 for details). Initialized with a 15-network parcellation as a prior, this method results in reliable within-individual estimates of networks by accounting for correlation variability both within an individual and between individuals. All network estimates for each individual were verified using seed-based region correlation to ensure the model-estimated networks were consistent with the underlying distributed correlation patterns (see Du et al. 2023).

### Anterior and Posterior Hippocampal Regions Constructed Within Individuals

Within-individual hippocampal regions were defined based on functional connectivity to the two cerebral networks targeted by Zheng et al. (2021) (Fig. 1). For each fixation run, the mean BOLD signal was extracted within the DN-A and SAL / PMN networks on the cerebral surface and correlated with the BOLD signal within each voxel of each individual’s hippocampal boundaries as identified by the automated FreeSurfer parcellation (Fischl et al. 2002, 2004). Every voxel along the hippocampus was assigned to the cerebral network with which it was most correlated. Unreliable voxels were excluded, including voxels along the edge of the hippocampal region, voxels with a temporal signal-to-noise ratio (SNR) below 50, low correlation voxels (*z*(*r*) < 0.03) and voxels with similar correlation values to both cerebral networks, defined as having a normalized correlation difference of less than 0.1 (|*z*(*r*)_DN-A_ – *z*(*r*)_SAL/PMN_/*z*(*r*)_DN-A_ + *z*(*r*)_SAL/PMN_| < 0.1). The SNR, low correlation, and ambiguity exclusions removed between 31.33% (P14) and 65.91% (P1) of voxels within the trimmed hippocampal mask.

**Figure 1.**
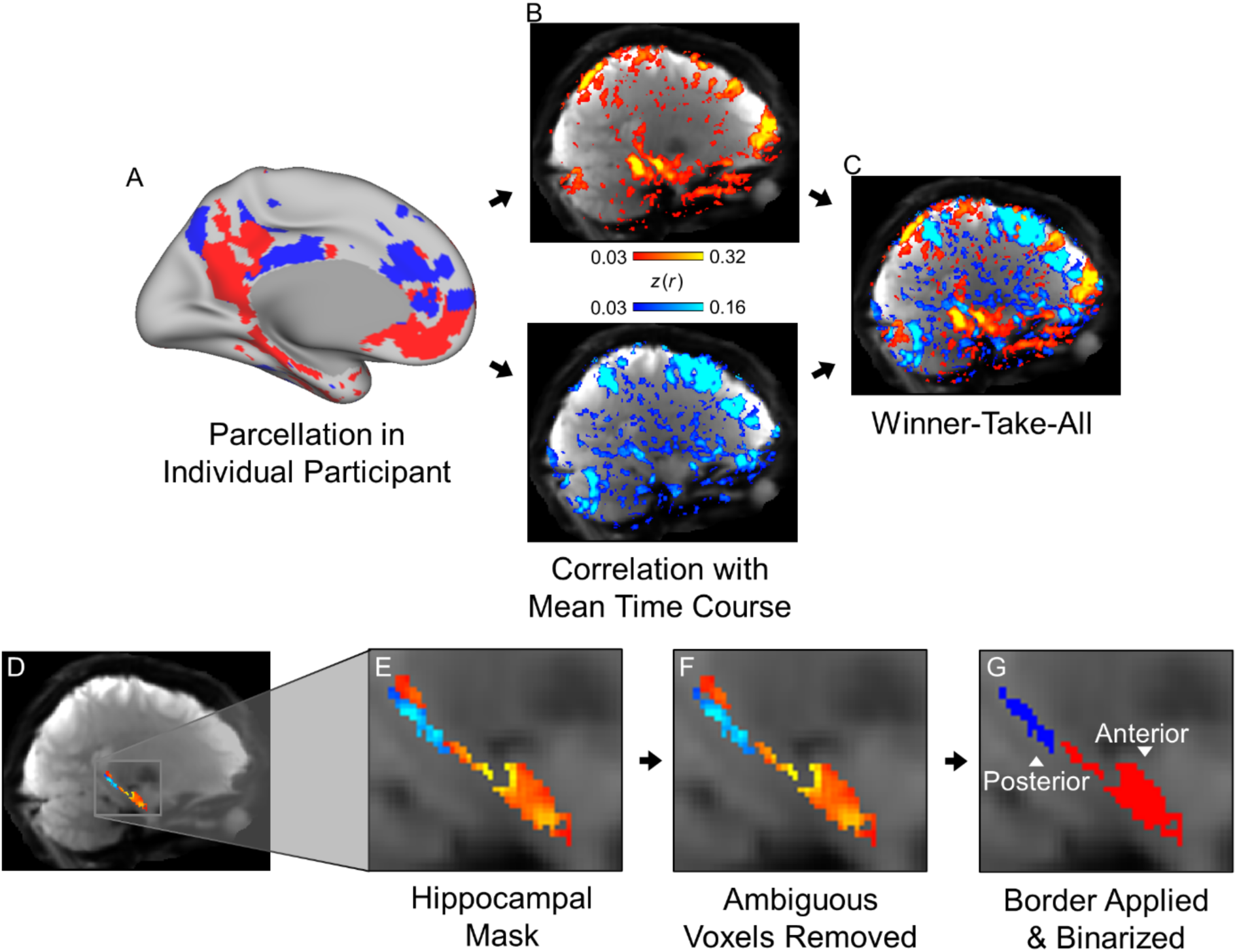
Posterior and anterior hippocampal regions are defined based on functional connectivity with cerebral networks. **(A)** In each participant, networks of interest (Default Network A, DN-A, red; Salience Network / Parietal Memory Network, SAL / PMN, blue) were identified on the cortical surface using a multi-session hierarchical Bayesian model (MS-HBM). **(B)** The mean BOLD time-course within each network’s boundaries was extracted and correlated with the BOLD time-course of every voxel in the brain volume. **(C)** Voxels in the volume were assigned to the network with which they were more correlated, and **(D, E)** masked for the hippocampus after excluding voxels with a signal-to-noise ratio below 50. **(F)** Within this hippocampal region, ambiguous voxels with comparable correlations to both networks (|*z*(*r*)_DN-A_ – *z*(*r*)_PMN_/*z*(*r*)_DN-A_ + *z*(*r*)_PMN_| < 0.1) were removed, and **(G)** a single network was assigned to define the anterior or posterior region. The network with > 50% of its assigned voxels anterior to border was chosen as the anterior-defining network, while the network with > 50% of assigned voxels posterior to the defined border was chosen as the posterior-defining network. In this representative participant, the border was defined at MNI coordinate Y = -31, and ∼84% of DN-A and SAL / PMN defined the anterior and posterior regions, respectively.

An MNI Y coordinate was chosen as the anterior / posterior border for each individual participant, and voxels assigned to each network were quantified posterior or anterior to that border. The border location was chosen such that the proportion of DN-A assigned voxels in the anterior region (relative to all hippocampal DN-A voxels) and proportion of SAL / PMN assigned voxels in the posterior region (relative to all hippocampal SAL/PMN voxels) would be equal, and vice-versa (DN-A proportion posterior to border equal to the SAL / PMN proportion anterior). In practice, this identified the coordinate where, moving along the long axis, one network’s assigned voxels began to be more prevalent while the other network’s assigned voxels began to be less prevalent. Following border assignment, hippocampal voxels anterior and posterior to the border were assigned to the defining network (DN-A or SAL / PMN) for whichever side of the border contained the majority of that network’s assigned voxels (voxels on the border were included in the anterior region). Voxels belonging to the non-assigned network were removed, and the remaining voxels were binarized to produce masks for the hippocampal regions that represented a best estimate of the anterior and posterior regions linked to the two separate cerebral networks.

The selectivity of these regions’ functional connectivity to the cerebral cortex was assessed by correlating the BOLD time course within each hippocampal region (whole brain regressed and bandpass filtered, as described above) with the BOLD time course of every vertex on the surface projected data (whole brain regressed, bandpass filtered, and smoothed to 2mm FWHM). The *z*-transformed correlation maps were visualized on the cerebral surface (Supplemental Figs. 2-3). Functional connectivity with *a priori* identified cerebral networks was quantified by averaging across all the vertices within each network (Fig. 2).

**Figure 2.**
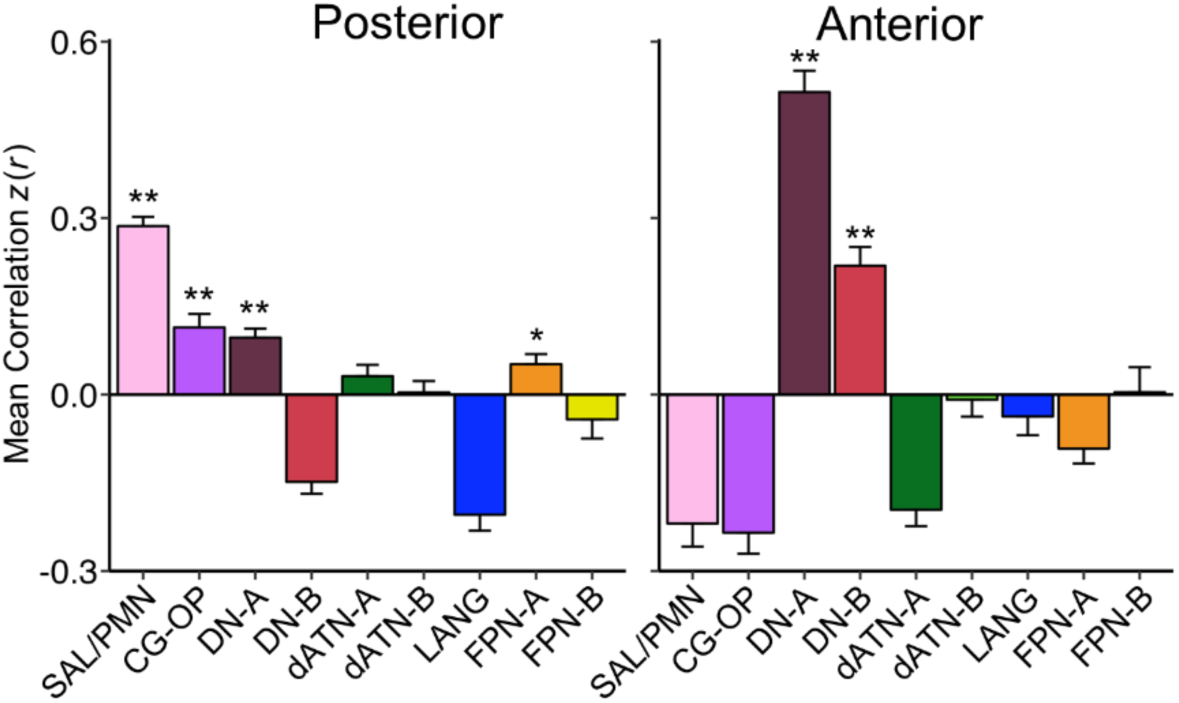
Posterior and anterior hippocampal regions are distinguished by their correlation patterns to distinct cerebral networks. Correlations between hippocampal regions and cerebral networks calculated within an individual were pooled across participants and averaged. In this pooled group average, the posterior hippocampal region was significantly correlated with SAL / PMN, CG-OP, and DN-A, with the strongest correlation to SAL / PMN. In contrast, the anterior hippocampal region was most strongly correlated with DN-A, with a significant but weaker correlation to DN-B. Correlations values are mean Fisher *z*-transformed Pearson’s *r* and error bars indicate the standard error around the mean *z*(*r*). Networks were clustered using a multi-session hierarchical Bayesian model (MS-HBM) and labeled according to the convention in Du et al. (2023). SAL / PMN = Salience Network / Parietal Memory Network; CG-OP = Cingulo-Opercular Network; DN-A = Default Network A; DN-B = Default Network B; dATN-A = Dorsal Attention Network A; dATN-B = Dorsal Attention Network B; LANG = Language Network; FPN-A = Frontoparietal Network A; and FPN-B = Frontoparietal Network B. Significance measured with a one-tailed t-test, H_o_: μ = 0, H_a_: μ > 0; ** = *p* < 0.001; * = *p* < 0.05

### Model-Free Seed-Region Based Functional Connectivity

A model free seed-based approach (e.g. Braga and Buckner 2017; Kosakowski et al. 2023) was used to assess the specificity of the functional connectivity to the hippocampus, rather than nearby cortical and sub-cortical structures. For this exploration, the pair-wise Pearson correlation was calculated between every voxel within an expanded hippocampal mask (59,343 – 68,354 voxels), based on an individual’s FreeSurfer parcellation, to every vertex on the cortical surface (81,924 vertices). The resulting correlation matrix was calculated for each resting-state fixation run, Fischer *z*-transformed, and averaged across all of an individual’s fixation runs to yield a single best estimate of that participant’s correlation structure. This correlation structure was then explored using Connectome Workbench’s wb_view software (Marcus et al. 2011; Glasser et al. 2013) by interactively choosing cerebral seed regions and visualizing the correlations from that seed region using the Jet look-up table (colorbar), excluding negative values. The correlations were thresholded to best see the functional connectivity topography on the cerebral surface and in the hippocampal volume (Supplemental Figs. 5-15).

### MRI Task Paradigms

Participants completed multiple runs of an Episodic Projection task (involving remembering and imagining the future), multiple runs of a Visual Oddball Detection task, and multiple runs of a Blocked Visual-Motor task. For additional details of the Episodic Projection and Visual Oddball Detection tasks, as well as additional acquired tasks not discussed here, see Du et al. (2023). The three tasks are briefly described below.

### Episodic Projection

The Episodic Projection task (10 runs, 10 min 17 sec each, 102 min 50 sec total) was an expanded version of the task from DiNicola et al. (2020) designed to encourage processes related to participants’ remembering their personal past and imagining their future. Target conditions asked participants to consider a brief written scenario about their past (Past Self) or possible future (Future Self) followed by answering a simple, multiple-choice question. Control conditions included general knowledge questions about the past (Past Non-Self) and future (Future Non-Self), as well as questions about the present (Present Self and Present Non-Self). The target vs. control contrasts of interest (Past Self vs. Present Self and Future Self vs. Present Self) have previously been shown to dissociate DN-A from DN-B (DiNicola et al. 2020), and detailed behavioral analyses suggest this dissociation may be driven by an increased reliance on mentally constructing scenes when answering target questions (DiNicola et al. 2023a; see also Hassabis and Maguire 2007). We utilized the Episodic Projection task here during scanning and also collected new behavioral ratings (see below) to replicate and further explore how the process of Scene Construction and other component processes modulate the present observed neural responses. Runs with more than two trials with no response from the participant were excluded. Additional conditions were acquired but not analyzed here.

### Visual Oddball Detection

The Visual Oddball Detection task (5 runs, 5 min 50 sec each, 29 min 10 sec total) assessed responses to infrequently-presented, task-relevant visual stimuli (similar to Wynn et al. 2015). Participants were presented stimuli sequentially in a rapid event-related design, with upper case “K”s and “O”s in either black or red, and asked to indicate whenever the target red upper case “K” was presented (10% of presentations). They pressed a key with their right index finger to indicate the presence of a red “K” and withheld their response for all other trials. Extended periods of passive fixation were included at the beginnings and ends of the run as an additional reference. The main portion of the task, analyzed here, was the extended, rapid event-related, continuous detection task. Individual runs were excluded from analysis if the participant missed more than 6 targets (20%).

### Blocked Visual-Motor

The Blocked Visual-Motor task (8 runs, 3 min 24 sec each, 27 min 12 sec total) required participants to view 30-sec blocks of visual flickering checkboards (4hz counter-phased black and white circular checkerboard). Seven or 8 times during each block, a pair of right and left red checks appeared unexpectedly and briefly for 0.25 sec. The participant’s task was to indicate by finger press whether the checks were near to the center or far from the center. Relevant to the present explorations, the visual-motor blocks were separated by multiple extended 18-sec blocks of passive visual fixation. This paradigm enabled exploration of transient responses at the transitions between blocks, including going from fixation to the visual-motor task and going from the visual-motor task to fixation. Individual runs were excluded from analysis if the participant got fewer than 80% of trials correct or failed to respond to 4 or more trials.

### Online Behavioral Task Paradigm

The goal of the online behavioral task was to assess the component processes that were embedded within each trial of the scanned Episodic Projection task, extending the procedure of DiNicola et al. (2023a). The idea motivating the approach is that each trial possesses an idiosyncratic set of features that shape how participants solve the trial, with differences from one trial to the next. Some trials may be more difficult than others, some may rely on scene construction more than others, etc. By assessing strategies used, on average, to complete each unique trial, the behavioral ratings of component processes can provide a means to explore what correlates with the MRI-measured functional responses.

The Episodic Projection task trials were administered to online participants via the Qualtrics survey tool (Qualtrics, Seattle, WA). Given the burden of doing the task and answering multiple strategy questions about each trial, data collection was spread across 10 separate cohorts of participants. As a control, 5 questions were repeated across every participant to test for cohort effects (no effects were noted), and 2 additional questions were included as attention checks (one probed whether participants were reading the questions, while another targeted task focus). To ensure only participants who fully engaged with the task were included, participants were excluded if they failed on measures of compliance including spending too little time on the survey, incorrect attention check responses, or stereotyped response patterns (as in DiNicola et al. 2023a). All exclusions were made prior to analyzing strategy composites. After exclusions, at least 25 participants contributed to strategy ratings for every trial.

Online presentation was similar to the scanned task. A few questions required minor wording changes to be applicable to online participants. After answering each individual question, participants reported the cognitive strategies they used when answering the question by giving a number from 1 (not used at all) to 7 (used extensively) for each of 21 strategies. Only after completing this reflection was the participant able to move forward and view the next trial.

Collected strategies included 13 from DiNicola et al. (2023a; adapted from Andrews-Hanna et al. 2010) and 8 additional strategies. To remain consistent with the functional characterizations described in DiNicola et al. (2023a), we did not include the 8 new strategies in the present analysis. The 13 utilized strategies probed the extent a given trial: (1) prompted the participant to think about their personal feelings (Pers_Feelings), (2) evoked emotions (Emotions), (3) asked the participant to rely on personal past experiences (Pers_Past_Exper), (4) led the participant to imagine a sequence of events unfolding (Sequence_Events), (5) led the participant to envision the locations of mentioned objects or places (Loc_Obj_Places), (6) led the participant to envision the physical locations of other people (Loc_People), (7) led the participant to speculate about others’ feelings (Others_Feelings), (8) made the participant consider general moral principles (Moral_Principles), (9) required the participant to think about their relationships to other people (Relationships), (10) made the participant consider the personality traits of other people (Others_Personality), (11) evoked visual imagery (Visual_Imagery), (12) was difficult (Difficulty), and (13) caused reliance on facts when answering (Facts).

### MRI Task Analysis

Run-specific general linear models (GLMs) were implemented through FSL version 5.0.4 first-level FEAT (Woolrich et al. 2001). Data were high-pass filtered with a cutoff at 100 sec to remove low frequency noise, and GLMs were then run for the cortical surface using whole-brain regressed data (smoothed to 2mm FWHM) and the entire MNI111-registered volume including the hippocampus (whole-brain regressed, not smoothed). Task contrasts were created separately for each run and averaged across runs to create an average task contrast using *fslmaths* (Smith et al. 2004). These mean values were extracted within each participant’s individualized cerebral networks (from the surface) and the anterior and posterior hippocampal regions (from the volume).

#### Episodic Projection

The Episodic Projection task was analyzed first at the level of conditions and then subsequently to isolate each individual trial separately. For the condition-level analysis, maps contrasted the Past Self vs. Present Self and Future Self vs. Present Self conditions. For the trial-level analysis, a GLM with separate regressors for every trial was employed (e.g., Hassabis et al. 2014; DiNicola et al. 2023a). The resulting single-trial parameter estimates were extracted and averaged across individuals. This allowed for extremely stable estimates of the BOLD response to be obtained for the condition contrasts and also for every unique trial question separately.

#### Visual Oddball Detection

The Visual Oddball Detection task was analyzed coding the target red “K”s and lures; other non-target stimuli were included in the implicit baseline. An additional GLM was run in the volume with cerebral DN-A activity as a regressor to control for the canonical task suppression effect that is known to affect the baseline of the hippocampus (Stark and Squire 2001). In a follow-up analysis to extract the time course of the response to oddball targets, the mean response to target stimuli in hippocampal regions was derived in each participant by first calculating the percent signal change in the cerebral DN-A network, anterior hippocampal region, and posterior hippocampal region during the task relative to periods of passive fixation at the beginning and end of the task, followed by regressing cerebral DN-A percent signal change from the hippocampal signal. The resulting adjusted activity was segmented into target presentation periods (from 2 TRs before presentation to 14 TRs after presentation), aligned, and averaged within individuals (excluding trials within 10 sec of the task onset cue to avoid interference). Finally, participants’ mean time courses were averaged to produce a group average impulse response time locked to the oddball targets.

#### Blocked Visual-Motor

Mean percent signal change within the hippocampal regions during the Blocked Visual-Motor task was calculated relative to fixation periods and then adjusted for DN-A network activity in each participant as described above. Signal was then averaged across participants and visualized for the entire extent of the run allowing the full evolution of the group averaged response to be visualized across block transitions.

### Behavioral Task Analysis

The mean trial-level strategy ratings from the online Episodic Projection task were *z*-scored and clustered (hclust function and ward.D2 amalgamation procedure in R) to create robust composite strategy scores for every Episodic Projection task trial. Strategies which clustered together in DiNicola et al. (2023a) were also closely clustered in this new data set, again yielding 5 clusters: (I) *Difficulty* (Facts and Difficulty strategies), (II) *Scene Constructio*n (Loc_Obj_Places and Visual_Imagery strategies), (III) *Others-Relevant* (Others_Feelings and Other_Personality strategies), (IV) *Self-Relevant* (Pers_Feelings and Emotions strategies), and (V) *Autobiographical* (Pers_Past_Experiences and Sequence_Events strategies; see Supplemental Fig. 16).

### Software and Statistical Analysis

Functional connectivity between brain regions was calculated in MATLAB (version 2019a; http://www.mathworks.com; MathWorks, Natick, MA) using Pearson’s product moment correlations. FreeSurfer v6.0.0, FSL, and AFNI were used during data processing. The estimates of networks in volume space were visualized in FreeView. The estimates of networks on the cortical surface were visualized in Connectome Workbench v1.3.2. Statistical analyses were conducted using R v4.2.1. Model-free seed-region confirmations were performed in Connectome Workbench v1.3.2. Network parcellation was performed using code from Kong et al. (2019) available on Github (https://github.com/ThomasYeoLab/CBIG/tree/master/stable_projects/brain_parcellation/Kong2019_MSHBM) with a 15-network prior (as in Du et al. 2023).

## Results

### The Anterior and Posterior Hippocampus Correlate with Distinct Cerebral Networks

Distinct patterns of functional connectivity were found in the hippocampus from the cerebral DN-A and SAL / PMN networks. Voxels in the anterior hippocampus were nearly all more strongly correlated with DN-A than SAL / PMN, while voxels more correlated with SAL / PMN were found in the posterior hippocampus (Supplemental Fig. 1). The effect was not absolute but was present in every participant and robust in most. Across participants the anterior-posterior border, where the relative differential connectivity to one network over the other flipped, fell between MNI Y coordinates -28 and -34, and in all cases DN-A was assigned to the region anterior to this border (henceforth referred to as the anterior hippocampal region) while SAL / PMN was assigned to the region posterior to the border (posterior hippocampal region) (Supplemental Fig. 1).

To explore whether the hippocampal regions were preferentially correlated with the targeted networks without imposing model assumptions, correlations from the two hippocampal regions were visualized across the entire cerebral surface. In all participants, visual inspection of the pattern of correlations from the hippocampal regions to the cerebral surface suggested preferential coupling to the two networks chosen to define each region (Supplemental Figs. 2 and 3). Quantification of correlations both aggregated across all 11 participants (Fig. 2) and examined at the individual level (Supplemental Fig. 4) confirmed the preferential connectivity of the hippocampal regions with the two networks.

Quantification of hippocampal-cortical correlations also revealed additional details. The posterior hippocampal region was most correlated with cerebral SAL / PMN (one-sided Student’s t-test, *t*(10) = 18, *p* < 0.001). Significant but weaker correlations were also observed to DN-A (*t*(10) = 6.25, *p* < 0.001) and the Cingulo-Opercular Network (CG-OP, *t*(10) = 4.92, *p* < 0.001). In contrast, the anterior hippocampal region was preferentially correlated with DN-A (*t*(10) = 14.2, *p* < 0.001) and to a lesser degree DN-B (*t*(10) = 6.78, *p* < 0.001), with no detectable coupling to SAL / PMN. The weaker, but still significant, correlations between the hippocampal regions and networks beyond the two networks initially chosen may suggest additional heterogeneity (see Reznik et al. 2023 for a recent analysis of DN-B in relation to correlations with the hippocampal formation). With these details in mind, the strong and preferential correlations to DN-A and SAL / PMN from the anterior and posterior hippocampal regions replicated Zheng et al. (2021) and provided a basis for detailed task-based functional explorations.

As a final control analysis to assess whether the estimates of the hippocampal functional connectivity patterns were affected by BOLD signal bleed from nearby cerebral cortex (i.e., entorhinal cortex, perirhinal cortex, and parahippocampal cortex) or thalamus, rather than signal within the hippocampus itself, we visualized correlations in the hippocampus from anteromedial and posteromedial cerebral cortical seed regions that reproduced the cerebral network patterns (Supplemental Figs. 5-15). Across all participants, both anterior and posterior seed regions in cerebral DN-A converged on correlation peaks within or near the hippocampus that were not continuations of immediately surrounding cerebral cortex. Seed regions within SAL / PMN produced similarly convergent correlations within the posterior hippocampus, distinct from nearby parahippocampal cortex or thalamus.

### The Anterior Hippocampus Responds During Remembering and Imagining the Future

During the Episodic Projection task, the anterior hippocampus displayed significant activity when individuals remembered their past (one-sided Student’s t-test, *t*(12) = 5.27, *p* < 0.001) or imagined their future (*t*(12) = 7.77 *p* < 0.001, Fig. 3A, top) relative to the control condition. In both contrasts the anterior hippocampus was significantly more active than the posterior hippocampus (two-sided Student’s paired t-test, Past: *t*(10) = 3.23, *p* < 0.01; Future: *t*(10) = 6.66, *p* < 0.001). This functional dissociation is also present within the broader cerebral networks associated with these hippocampal regions (DN-A and SAL / PMN, Fig. 3A bottom), replicating the specialization of cerebral DN-A for remembering the past and imagining the future (DiNicola et al. 2020, 2023b; Du et al. 2023).

**Figure 3.**
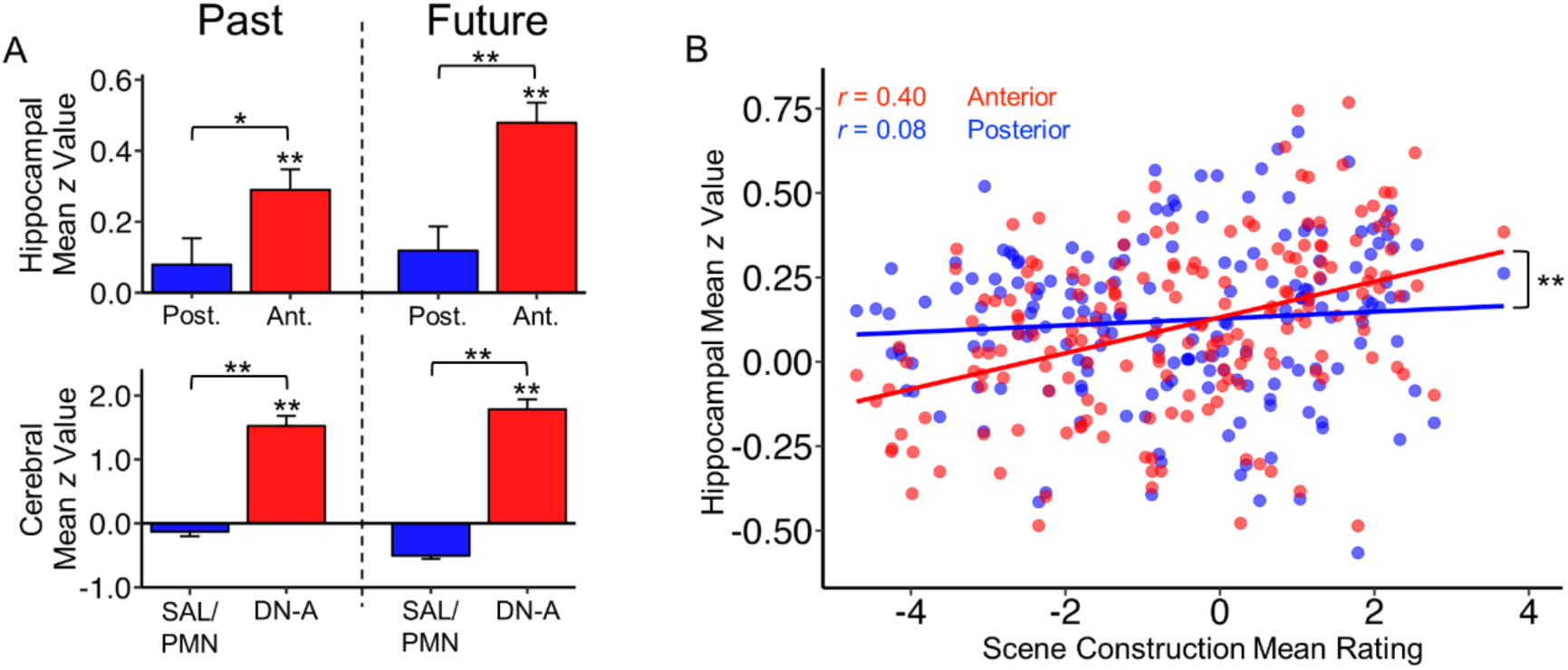
The anterior hippocampus responds during remembering and imagining the future and tracks scene construction. **(A)** Condition-based analysis of the Episodic Projection tasks reveal (top) increased activity within the anterior hippocampal region, but not the posterior hippocampal region, during contrasts targeting remembering and imagining the future (one-tailed t-test, H_o_: μ = 0, H_a_: μ > 0; Past = Past Self – Present Self; Future = Future Self – Present Self). The cerebral DN-A network (below) shows the same pattern, highlighting the similar functional behavior of cerebral DN-A and the anterior hippocampus. **(B)** A trial-level behavioral breakdown of activity within the anterior and posterior hippocampal regions reveals that activity within the anterior hippocampal region is correlated with the behavioral scene construction scores (*p* < 0.001), while activity in the posterior hippocampus is not (*p* = 0.27). Values are mean Fisher *z*-transformed beta values averaged across all participants. Error bars display the standard error of the mean. Paired t-tests are two tailed (H_o_: μ_1_ = μ_2_, H_a_: μ_1_ ≠ μ_2_). ** = *p* < 0.001; * = *p* < 0.05

### The Response in the Anterior Hippocampus Tracks Scene Construction

The response of the anterior hippocampus across Episodic Projection task trials was significantly correlated with the behavioral estimates of scene construction (Fig. 3B; Pearson’s correlation, *r* = 0.40, 95% CI [0.27 0.52], *t*(178) = 5.82, *p* < 0.001), while the posterior hippocampus showed minimal correlation (Fig 3B; *r* = 0.08, 95% CI [-0.06 0.23], *t*(178) = 1.12, *p* = 0.27). Direct contrast of the anterior versus posterior hippocampal regions revealed a significant region * scene construction interaction (linear mixed effects model, type II Wald F test, *F*(1,178) = 36.537, *p* < 0.001). Control analyses, testing previously-described relations between cerebral networks and behavioral estimates, verified that the novel trial-level strategy estimates collected here behave consistently with previously-reported results (Supplemental Figs. 16 and 17). Thus, activity within the anterior hippocampus tracks scene construction and again differentiates its response properties from the posterior hippocampus.

In their report, Zheng et al. (2021) postulated that the anterior hippocampal region may be involved in self-relevant processes. The above results suggest a component process driving the anterior hippocampus response in our episodic remembering task was scene construction (as hypothesized by Hassabis and Maguire 2009). However, that does not mean the region’s response cannot also track component processes related to self-relevance, perhaps reflecting the multiple cerebral networks that are linked to the anterior hippocampal region (e.g., DN-B; Fig. 2). To explore this possibility, we tested the correlation of our behavioral estimate of self-relevance with the anterior hippocampus and found a weaker but still significant correlation (Supplemental Fig. 18, Pearson’s correlation, *r* = 0.17, 95% CI [0.02 – 0.31], *t*(178) = 2.26, *p* < 0.05). A multiple regression linear model including both scene construction and self-relevance behavioral composites as predictors significantly predicted activity in the anterior hippocampal region (*F*(2, 177) = 20.2, *p* < 0.001), and further showed that scene construction accounted for more variance in the anterior hippocampal region BOLD signal (Adjusted *R^2^*_Full Model_ = 0.18, *p*_Scene Construction_ < 0.001, *p*_Self Relevance_ < 0.05; *R^2^*_Scene Construction_ = 0.16; *R^2^*_Self Relevance_ = 0.02). Thus, we do find some evidence that the anterior hippocampal region tracks self-relevance, but the major association linked to its activity was scene construction.

To broadly assess the relative contribution of scene construction versus other component cognitive processes to anterior hippocampal activity, we controlled for trial difficulty and fit an additional multiple regression linear model including behavioral composite scores for scene construction and self-relevance, as before, as well as a composite scores for autobiographical features (considering personal past experiences and considering a sequence of events). This difficulty-corrected model continued to significantly predict activity within the anterior hippocampus (*F*(3, 176) = 10.81, *p* < 0.001), with the scene construction composite as the only significant predictor (Adjusted *R^2^*_Full Model_ = 0.14, *p*_Scene Construction_ < 0.001, *p*_Autobiographical_ = 0.13, *p*_Self Relevance_ = 0.86).

Finally, as an extreme exploration of the hypothesis that the anterior hippocampus responds to the process of scene construction, we restricted our analyses to only control condition trials (Present Self, Past Non-Self, Future Non-Self, and Present Non-Self). Even in these trials, designed to minimize episodic memory demands, activity within the anterior hippocampus was significantly correlated with the behavioral estimates of scene construction (Pearson’s correlation, *r* = 0.21, 95% CI [0.03 – 0.38], *t*(118) = 2.36, *p* < 0.05), and was significantly more sensitive to scene construction than the posterior hippocampus (posterior hippocampus: *r* = 0.00, 95% CI [-0.18 – 0.18], *t*(118) = 0.04, *p* = 0.97; hippocampal region * scene construction interaction: linear mixed effects model, type II Wald F test, *F*(1, 118) = 10.7, *p* < 0.01).

### The Posterior Hippocampus Transiently Responds During Oddball Detection

We tested the hypothesis that the function of the posterior hippocampus might be understood by anchoring from the role of the SAL / PMN in detecting salient, novel events and responding to task transitions (Konishi et al. 2001; Fox et al. 2005; Dosenbach et al. 2006; Seeley et al. 2007; Seeley 2019). We first explored the response of the hippocampal region in a Visual Oddball Detection task (requiring detection of red Ks among rapidly-presented letters). In the cerebral cortex, the set of regions activated by the salient targets included the SAL / PMN network (Fig. 4A; Supplemental Fig. 19). Quantification of this response (Fig. 4B) confirmed that activity to salient targets was primarily within the cerebral SAL / PMN (one-sided Student’s t-test, *t*(10) = 5.83, *p* < 0.001) and closely related CG-OP (*t*(10) = 9.92, *p* < 0.001) networks. Consistent with the dissociation raised in Zheng et al. (2021), we also observed large decreases in response to salient targets in both the DN-A and DN-B networks (Fig. 4B; see also Du et al. 2023). Given these results for the cerebral cortex, we turned our attention to the hippocampal regions, where we observed a clear functional dissociation.

**Figure 4.**
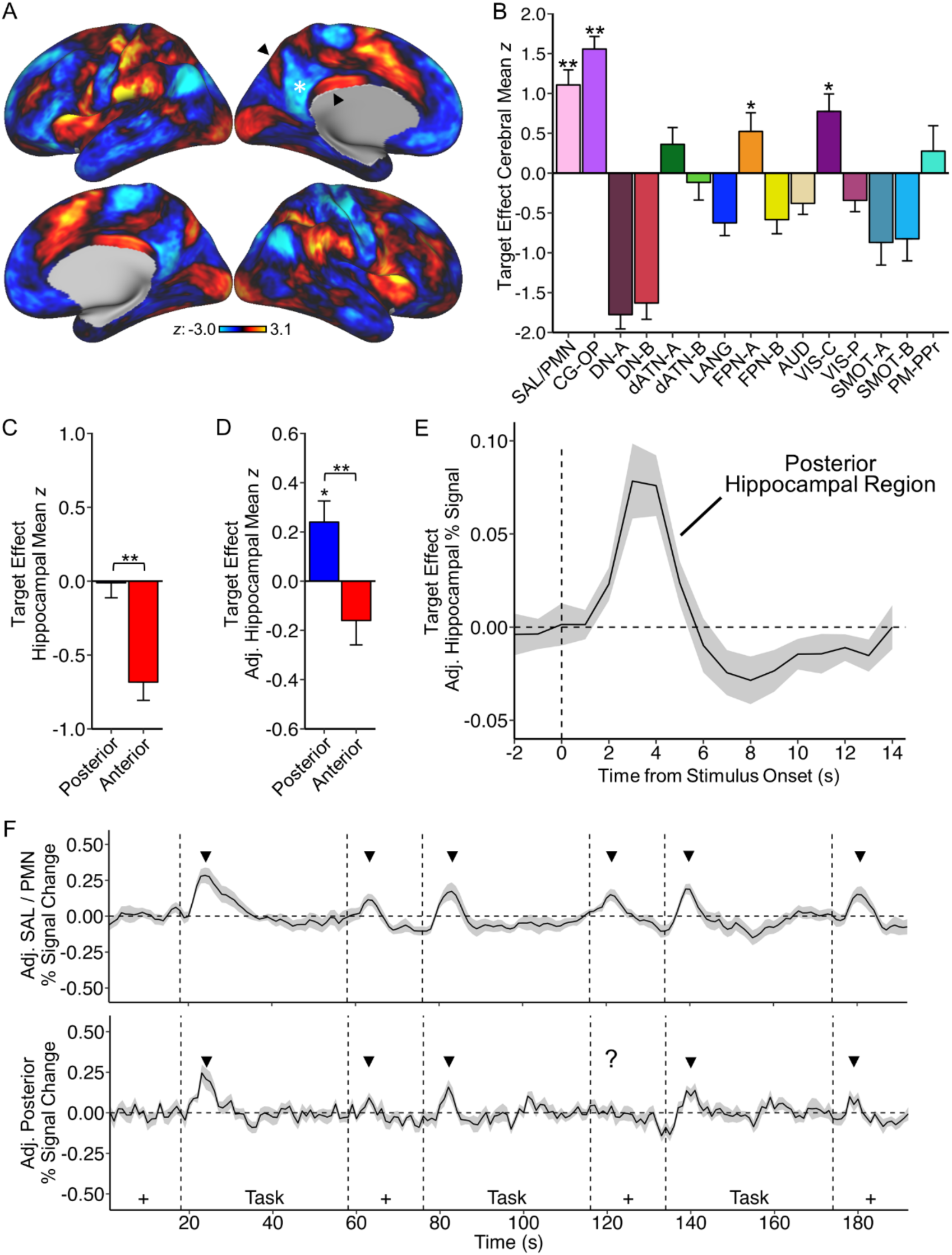
The posterior hippocampus transiently responds to oddball targets and task transitions. **(A)** Group-averaged activity on the cerebral surface in response to visual oddball targets reveals a robust positive response in regions of the SAL / PMN network including the posteromedial cortex (indicated by arrows). In contrast, nearby regions of the canonical Default Network show decreased activity (indicated by asterisk). **(B)** Quantification of this response by cerebral network reveals significant increases in the SAL / PMN and related CG-OP networks. DN-A and DN-B display prominent activity decreases. **(C)** Within the hippocampus, the posterior region is significantly more active than the anterior region, although both display relatively low activity (meaning the reference baseline is high). **(D)** Regressing DN-A activity from both hippocampal regions shifts the baseline. **(E)** Plotting the mean percent signal change within the posterior hippocampal region time locked to the target oddball events shows a transient, canonical hemodynamic response. The dashed line shows the onset time of the targets. The shading around the response indicates the standard error of the mean. **(F)** Adjusted time courses of the Blocked Visual-Motor task are shown for the cerebral SAL / PMN network (top) and for the posterior hippocampal region (bottom). The dashed lines indicate the transitions between blocks with the notation at the bottom indicating the block types (Task or fixation, +). The arrows indicate the transient responses at the block transitions. Significance statistics are either a one-tailed t-test for each hippocampal region (H_o_: μ = 0, H_a_: μ > 0), or a two-tailed paired t-test to assess for differences between the hippocampal regions (H_o_: μ_1_ = μ_2_, H_a_: μ_1_ ≠ μ_2_). Error bars indicate standard error of the mean. Additional network abbreviations: AUD = Auditory; VIS-C = Visual-Central; VIS-P = Visual-Peripheral; SMOT-A = Somatomotor A; SMOT-B = Somatomotor B; PM = Premotor; PPr = Posterior Parietal Rostral. ** = *p* < 0.001; * = *p* < 0.05

The posterior hippocampal region responded to salient oddball targets significantly more than the anterior hippocampal region (Fig. 4C; two-sided paired Student’s t-test, *t*(10) = 5.84, *p* < 0.001). A feature of this differential response is a relatively high baseline within the hippocampus, shifting the overall responses (see Stark and Squire 2001). To better visualize the relative differential response between the hippocampal regions, we regressed DN-A activity, which also showed the canonical “deactivation” or task suppression effect. After regression, the posterior hippocampal region displayed significant positive activation against the new baseline (Fig. 4D; one-sided Student’s t-test, *t*(10) = 2.8, *p* < 0.05), while also remaining significantly more active than the anterior hippocampus (*t*(10) = 4.64, *p* < 0.001). This *post-hoc* analysis should be thought of as helping to see a pattern rather than changing a pattern, as the relative differential response between the anterior and posterior hippocampal regions is the same and significant in both analyses (it is the *relative* baseline that shifts).

As a final analysis to investigate the posterior hippocampal response, we averaged activity within the posterior hippocampal region after each of the target trials to visualize the evolution of the hemodynamic response. A robust, canonical transient hemodynamic response was observed time-locked to the oddball targets (Fig. 4E). These results indicate that the posterior hippocampus responds transiently to simple salient target stimuli.

### The Posterior Hippocampus Transiently Responds at Task Block Transitions

The observation that the posterior hippocampus responds to visual oddball targets, while predicted by the correlation of the posterior hippocampus with the cerebral SAL / PMN network, was nonetheless surprising given there were no declarative or associative memory demands. To generalize this effect to another independent paradigm, we explored anterior and posterior hippocampal region responses in a Blocked Visual-Motor task paradigm that possessed embedded transitions at the beginnings and endings of the task blocks (e.g., Konishi et al. 2001; Fox et al. 2005; Dosenbach et al. 2006).

Fig. 4F illustrates the results. At each block transition – including both transitions from visual fixation to the active task block and the reverse transition going from the active block to passive fixation – there was a transient response observed in the posterior hippocampus that paralleled the block transition effect in the cerebral SAL / PMN network. Supplementary Fig. 20 shows results both with and without regression of cerebral DN-A activity. While the response was largest in the initial transition from fixation to the active task block, it was clearly present in transitions to the passive fixation blocks, where minimal dynamic task demands are present after the transition. These results support that the posterior hippocampus responds transiently at task block transitions.

## Discussion

The anterior hippocampal region, like its partner cerebral network DN-A, responds during remembering and imagining the future, tracking the process of scene construction – a core component of episodic remembering linked to hippocampal function (Hassabis and Maguire 2007, 2009). By contrast, the posterior hippocampal region, like its partner cerebral network SAL / PMN, responds transiently during oddball detection and also at the transitions between task blocks. The response of the posterior hippocampus to oddball targets and task transitions is present despite the absence of declarative or associative memory demands. We discuss implications of these findings and limitations for understanding specialization of the long axis of the hippocampus.

### The Anterior Hippocampus Tracks Scene Construction During Remembering and Imagining the Future

Zheng and colleagues (2021) dissociated the anterior and posterior hippocampus based on the regions’ functional connectivity with cerebral networks. The present results replicate their observations and add further details. Our estimates, like those of Zheng et al. (2021), reveal that the anterior hippocampus is correlated with the distributed cerebral network historically referred to as the “Default Network”. Based on recent within-individual precision estimates, we examined the specific relation to DN-A, a cerebral network involved in episodic remembering, separately from the distinct parallel cerebral network, DN-B, specialized for social inferences (e.g. Braga and Buckner 2017; Braga et al. 2019; DiNicola et al. 2020, 2023a; Deen and Freiwald 2022; Du et al. 2023; Edmonds et al. 2023). The anterior hippocampus was coupled to both DN-A and DN-B, with the correlation to DN-A being substantially greater (Fig. 2), leading to our prediction that the anterior hippocampus might track the process of scene construction (Hassabis and Maguire 2007, 2009; DiNicola et al. 2023a). This prediction was borne out.

The anterior hippocampus was significantly active during contrasts targeting remembering the past and imagining the future, and more so in both condition contrasts than the posterior hippocampus (Fig. 3A). Note that these contrasts are designed to minimize contributions of self-relevance by subtracting that component from the baseline condition (see the clear double dissociation of cerebral networks DN-A and DN-B in Du et al. 2023, independently replicating data in DiNicola et al. 2020). Moreover, the response in the anterior hippocampal region across trials significantly correlated with the component process of scene construction (Fig. 3B). This relation with scene construction was absent for the posterior hippocampal region. Furthermore, within the anterior hippocampal region, the relation with scene construction was present in the control trials (Supplementary Fig. 18). Thus, by all analyses, converging with our previous observations about the cerebral DN-A network (DiNicola et al. 2023a) and observations by others (Hassabis and Maguire 2007, 2009), the anterior hippocampus tracks the component process of scene construction.

The positive evidence presented here for a role of the anterior hippocampus in scene construction should not be taken to mean the region only responds to such processes. Detailed analyses of the correlation of the anterior hippocampal region to the cortex also revealed evidence for linkage to the DN-B cerebral network (Fig. 2), including in many of the individual participants (Supplemental Fig. 4). Reznik and colleagues (2023), using high-field MRI, recently demonstrated that anterior regions of the hippocampal formation, particularly a region within the entorhinal cortex, was linked to the cerebral DN-B network. They further revealed a clear separation between DN-A and DN-B medial temporal lobe linked regions (see Reznik et al. 2023 their Figs. 3 and 5). All these results converge to suggest that the long axis of the human hippocampus is not solely defined by interactions with cerebral networks DN-A and SAL / PMN, but rather that there are certainly additional networks that also link to the hippocampal formation. The correlation with DN-A is robust and perhaps the most dominant contribution to the anterior hippocampus, driving the functional response properties we observed here in our analyses. Future explorations at high resolution that combine functional connectivity and task-based manipulations are warranted to dig deeper, especially in relation to processing within the social domain linked to the DN-B cerebral network.

### The Posterior Hippocampus Responds to Salient Transients

A clear but surprising finding of the present paper is the transient response in the posterior hippocampus to oddball targets and transitions between task blocks (Fig. 4). The motivation to explore such processes was drawn directly from the network estimated to be linked to the posterior hippocampus. In this regard, our interpretation of the network correlation pattern for the posterior hippocampal region differs from that of Zheng and colleagues (2021). We hypothesize the major network correlated with the posterior hippocampal region is the same as the network described by Seeley and colleagues as the Salience Network (SAL; Seeley et al. 2007; Seeley 2019; see also Dosenbach et al. 2006), with weaker coupling to the nearby and closely related Cingulo-Opercular Network (CP-OP; Dosenbach et al. 2006; Seeley 2019). In contrast, Zheng and colleagues proposed that the posterior hippocampal network is linked to memory functions through a sensitivity to a familiarity response. The basis of their hypothesis is prior studies that have demonstrated regional responses at and around the PMN to item repetitions in memory paradigms (e.g., Gilmore et al. 2015; see also Nelson et al. 2013; Chen et al. 2017). Our alternative interpretation, based on the observation that the primary posterior hippocampal network is spatially similar to the SAL network (see Du et al. 2023 for discussion), led us down a different path. We hypothesized and found evidence that the posterior hippocampus responds to oddball targets and task transitions.

These findings are consistent with work describing a role for the hippocampus in processing surprise or novelty (e.g., Rolls et al. 1989; Knight 1996; Fyhn et al. 2002; Kumaran and Maguire 2009). Transient responses were noted to isolated, repeating simple letter stimuli when they were oddball targets and at task block transitions, including when the transition was away from an active task and toward passive fixation. Previous neuroimaging investigations have noted transient responses in the hippocampus at the offset of complex, meaningful stimuli hypothesized to support consolidation (e.g., Ben-Yakov and Dudai 2011; Barnett et al. 2022). Here, transient responses were observed in paradigms with no obvious declarative or associative memory demands or meaningful materials to be encoded – just salient stimuli and transitions that required a task set change or orienting response. Our results connect the work on hippocampal novelty responses to the literature on cerebral salience processing (e.g., Dosenbach et al. 2006; Seeley et al. 2007; Seeley 2019) and motivate further study of the posterior hippocampus in this context. It is an open question whether the transient responses found here reflect a mechanism that is critical to memory or whether they reveal a processing role of the hippocampus that might best be conceived outside of the traditional focus on declarative memory.

### Limitations and Future Directions

Despite the improvements made possible by adopting a within-individual, network-based approach, our exploration of the hippocampus using fMRI was still limited by several technical considerations. First among these is signal dropout in the most anterior portions of the hippocampus and surrounding cerebral regions. Beyond preventing the inclusion of these regions in our analyses (via screening for tSNR), the lack of signal in regions of the anterior parahippocampal gyrus makes it difficult to assess the possibility of signal bleed from this portion of cortex into included regions of the anterior hippocampus. It is possible that anterior parahippocampal or entorhinal cortex activity, which we are unable to reliably measure in our study, is contributing to responses within the anterior hippocampus. In addition to dropout, the resolution of our BOLD acquisition (2.4 mm isotropic) becomes a notable hindrance when studying a structure as small as the hippocampus, resulting in ambiguity around the spatial location of activations and correlations (see Supplemental Figs. 5-15). Future work will benefit from acquisitions at high field with higher resolution to address these issues as recently conducted (Kwon et al. 2023; Reznik et al. 2023).

A future direction is to directly explore the relation of the posterior hippocampal transient responses observed here with memory repetition effects that have been the emphasis of prior work on the cerebral SAL / PMN network (and a proposed processing function of the posterior hippocampus; Gilmore et al. 2015; Chen et al. 2017; Zheng et al. 2021; Kwon et al. 2023; see also Gilmore et al. 2019). It is notable that maps of repetition effects can look remarkably similar to the visual oddball detection effects observed here. For example, the individual participant maps of repetition effects in Kwon et al. (2023; their Figs. 8, see S3 and S8) are highly similar to the individual participant oddball detection effect maps in the present work (Supplemental Fig. 19, see P3 and P6).

There are multiple possible ways the two sets of findings might relate. One possibility is that repeating targets in certain forms of recognition or incidental memory paradigms might make the repeated targets more salient. Response times tend to be faster to old items in old / new recognition decisions and confidence goes up with the number of repetitions. Thus, one avenue for future exploration is to explore old / new recognition memory and item repetitions in classic paradigms and ones that explicitly shift the relevance of the repetition (e.g., press only to old items or press only to new items; Shannon and Buckner 2004). More broadly, it will be interesting to explore processing models that link novelty and detection of salience to memory.

### Conclusions

The hippocampus can be robustly dissociated along the long axis based on connectivity to cerebral networks. The anterior hippocampus is correlated with the cerebral network DN-A, while the posterior hippocampus is correlated with the cerebral network SAL / PMN. The network-defined anterior and posterior regions display a functional double dissociation, highlighting the value of network-driven, domain-agnostic explorations of hippocampal function. The anterior hippocampus is sensitive to remembering, tracking the component process of scene construction; the posterior hippocampus responds transiently to salient stimuli and task transitions.

## Acknowledgments

We thank the Harvard Center for Brain Science neuroimaging core and FAS Division of Research Computing for providing critical support, specifically T. O’Keefe for assisting with data processing optimization and R. Mair for MRI physics support. We would also like to thank J. Du, H. Kosakowski, W. Sun and M. Elliot for their valuable input and discussions. K. Ntoh and F. Davy-Falconi assisted with data acquisition. The multi-band EPI sequence was generously provided by the Center for Magnetic Resonance Research (CMRR) at the University of Minnesota, and E. Fedorenko, T. Konkle, and R. Saxe generously provided task stimuli. P.A.A. thanks Daniel Schacter, Rick Born, and Mark Andermann for valuable insights as this work evolved.

## Grants

Supported by NIH grant MH124004, NIH Shared Instrumentation grant S10OD020039, and NSF grant 2024462. L.M.D. was supported by NSF Graduate Research Fellowship Program grant DGE1745303. Any opinions, findings, and conclusions or recommendations expressed in this material are those of the authors and do not necessarily reflect the views of the National Science Foundation.

## Supplemental Figure Legends

**Supplemental Figure 1.**
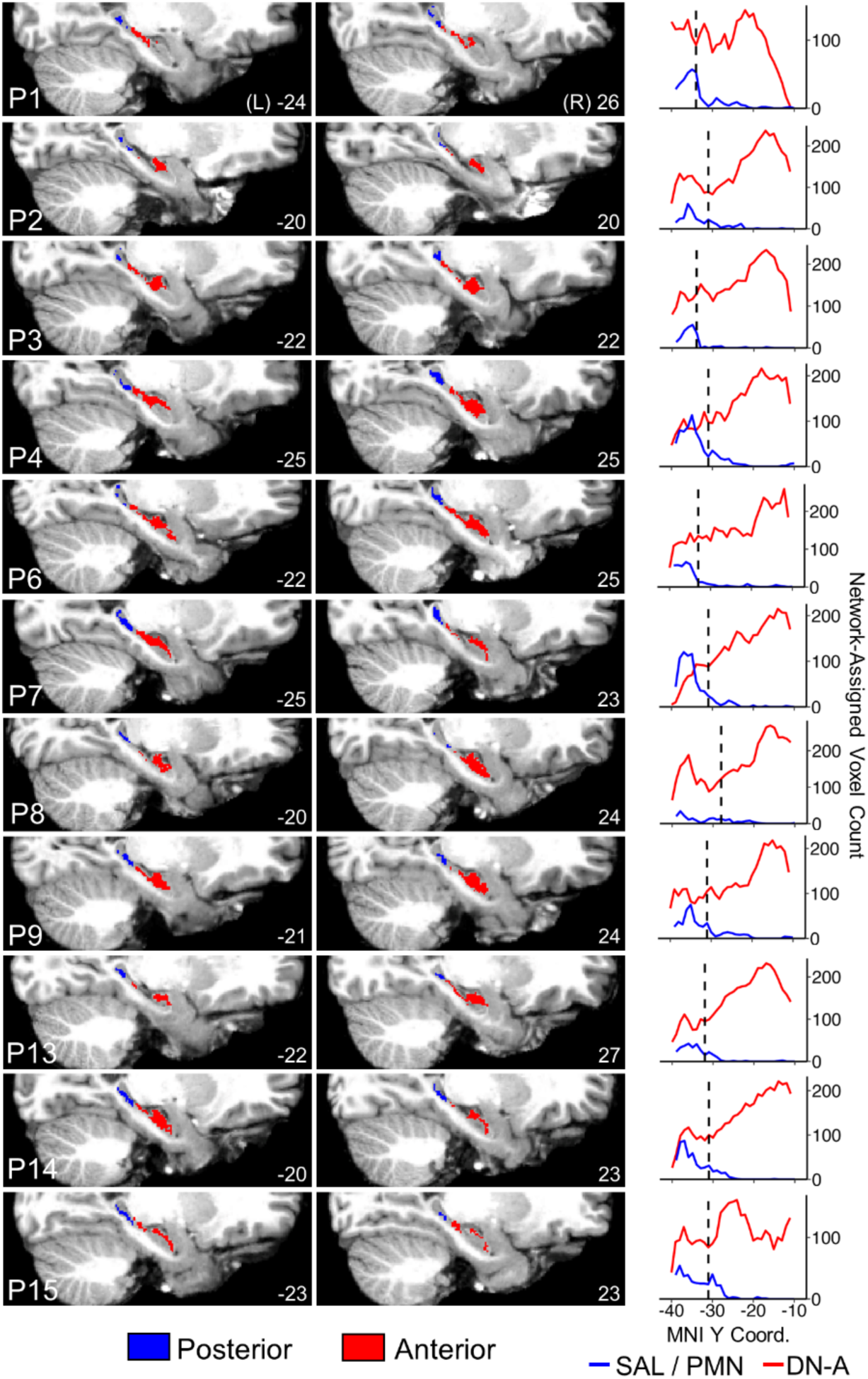
Posterior and anterior hippocampal region assignments to SAL / PMN and DN-A are consistent across participants. For each participant a single representative sagittal slice for the left (L) and right (R) hemisphere is shown, along with a plot of SAL / PMN and DNA assigned voxel counts along the hippocampal long axis. The MNI X coordinate for each slice is given in the bottom right of each image, and the location of the anterior / posterior border is shown by the black dashed line in the plots on the right. In all participants, SAL / PMN was the appropriate choice for defining the posterior hippocampal region (blue), while DN-A was appropriate for defining the anterior hippocampal region (red). While the size of the anterior and posterior regions varies between individuals, SAL / PMN correlated voxels were consistently present in the posterior hippocampus, while DN-A correlated voxels were the majority of voxels in the anterior hippocampus. The anterior / posterior border defined for each participant were, from top to bottom, MNI Y = -34, -31, -34, -31, -33, -31, -28, -31, -33, -32, -31, -31.

**Supplemental Figures 2 and 3.**
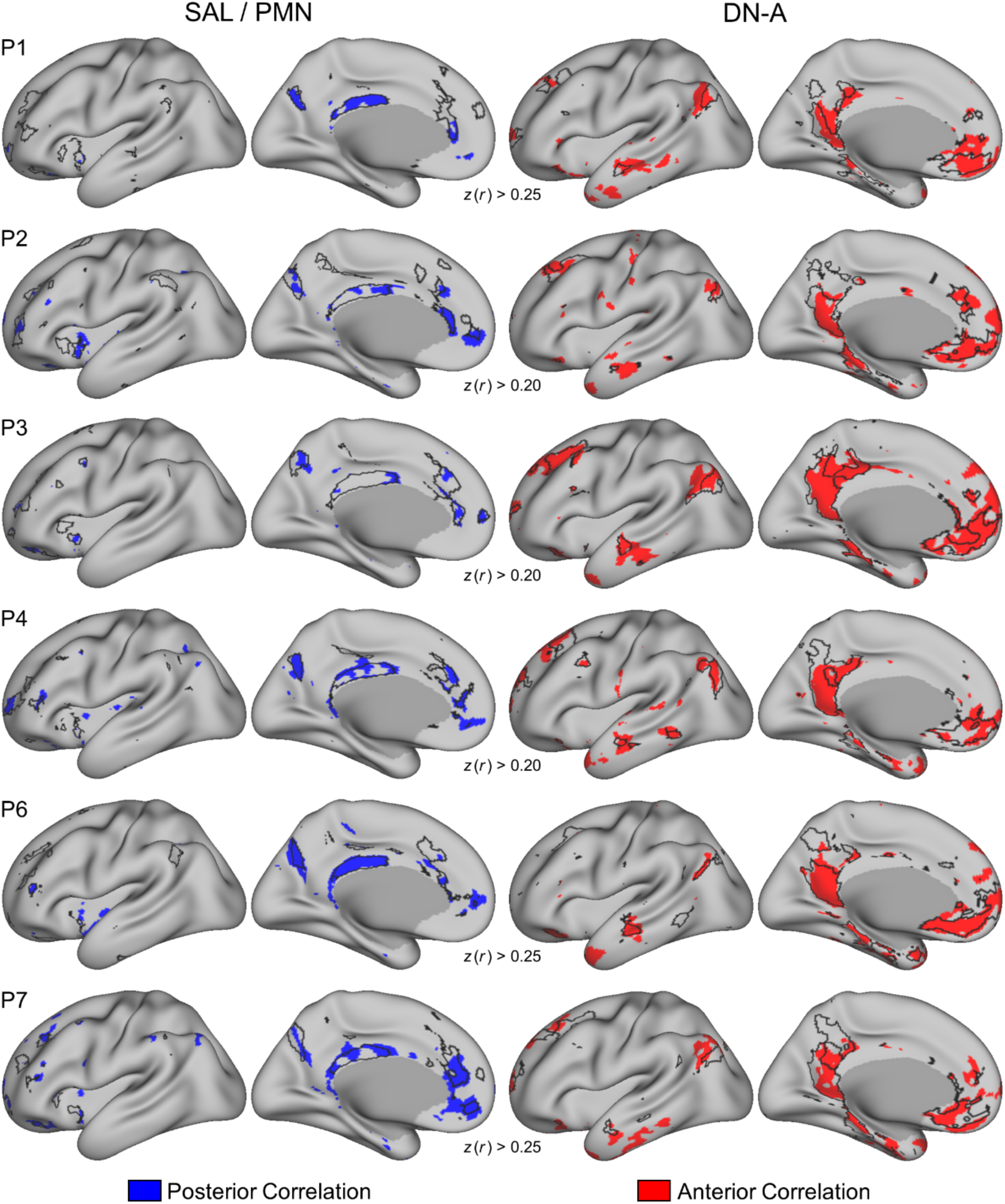
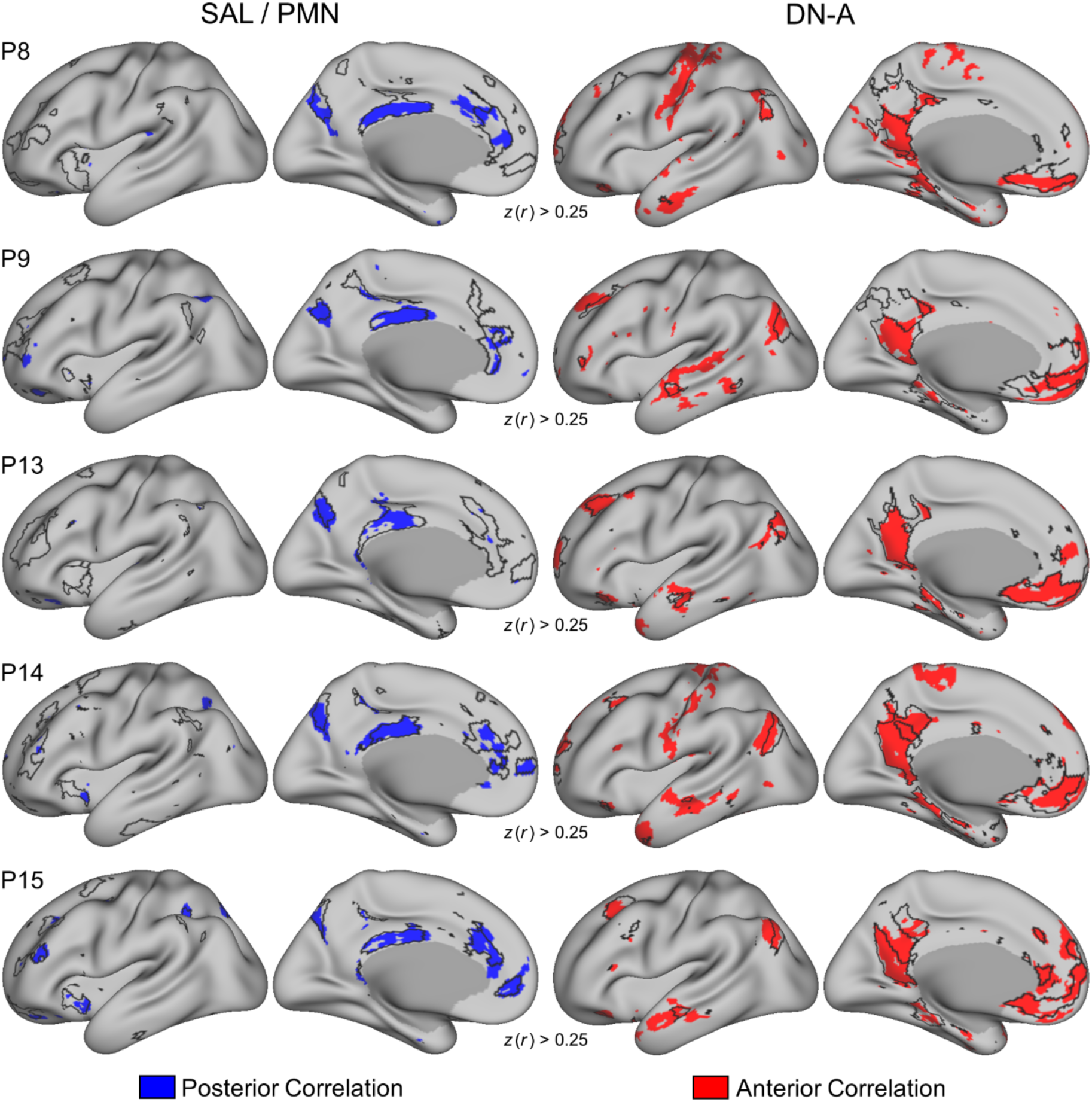
Functional connectivity from the posterior and anterior hippocampal regions recapitulates diagnostic features of the cerebral SAL / PMN and DN-A networks. Correlations to the cerebral surface from the individual-specific hippocampal regions in all participants followed the topography of the networks used to identify each region (SAL / PMN for the posterior region, DN-A for the anterior region). **(Left)** Correlations from an individual’s posterior hippocampal region (blue) along with their multi-session hierarchical Bayesian model (MS-HBM) derived cerebral SAL / PMN network outline. Notable regions within the SAL / PMN network include posteromedial and posterior cingulate cortices, as well as the anterior insula. **(Right)** Correlations from an individual’s anterior hippocampal region (red) along with their MS-HBM derived cerebral DN-A network outline. Notable regions within the DN-A network include retrosplenial, parahippocampal, and dorsolateral prefrontal cortices. Correlations displayed as Fisher *z*-transformed Pearson’s *r*, thresholded at the *z*-value indicated below each participant’s maps.

**Supplemental Figure 4.**
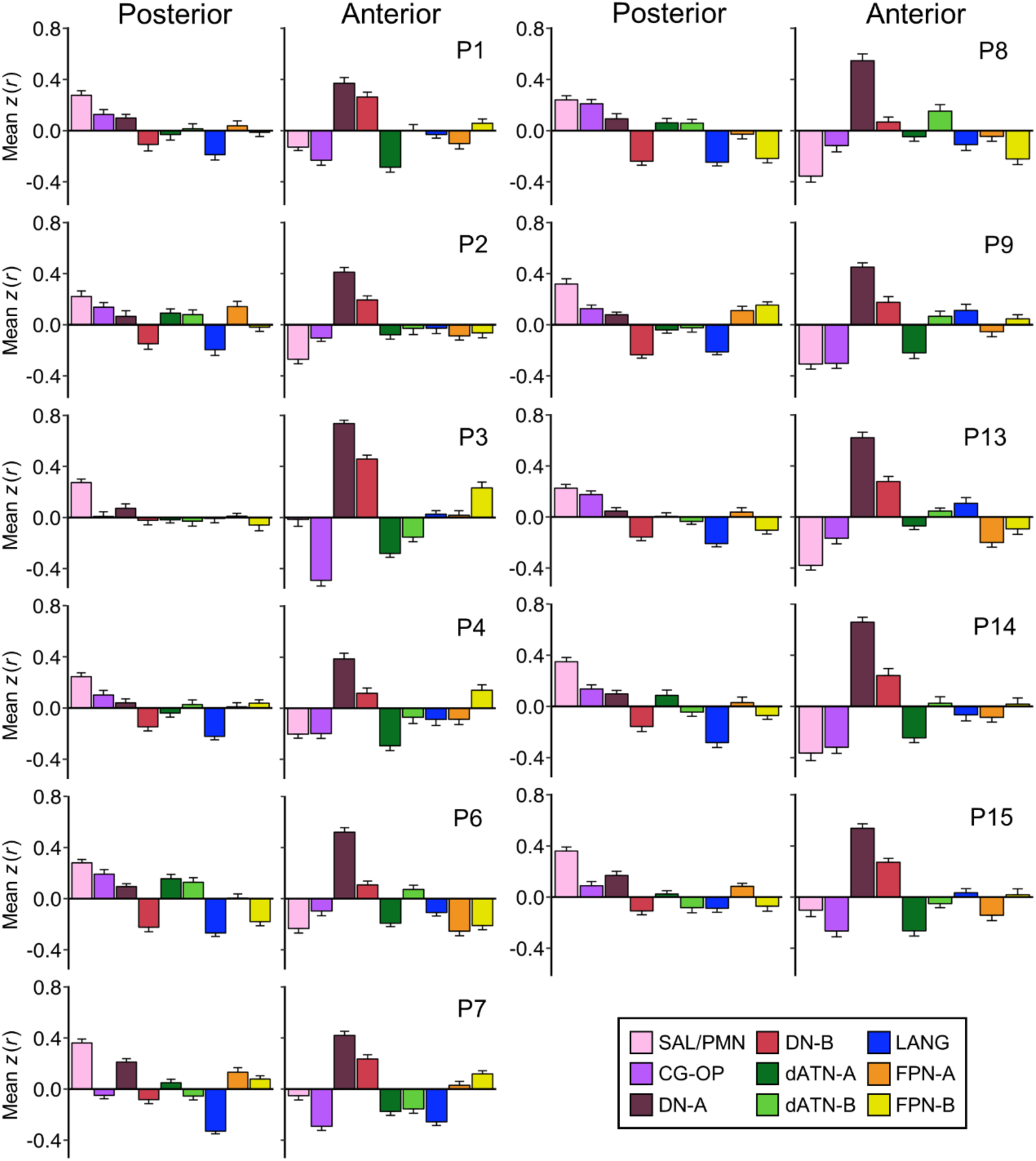
Individual-specific posterior and anterior hippocampal regions are preferentially correlated with the SAL / PMN and DN-A cerebral networks. Quantification of the mean correlations from individual participant’s posterior and anterior hippocampal regions to cerebral networks. In all participants the posterior hippocampal region was most correlated with cerebral SAL / PMN. The anterior hippocampal region was most correlated with DN-A, with many participants also showing high correlations with DN-B. Correlation values are mean Fisher *z*-transformed Pearson’s *r*, and error bars indicate the standard error around the mean *z*(*r*) across separate fixation scan runs.

**Supplemental Figures 5 to 15.**
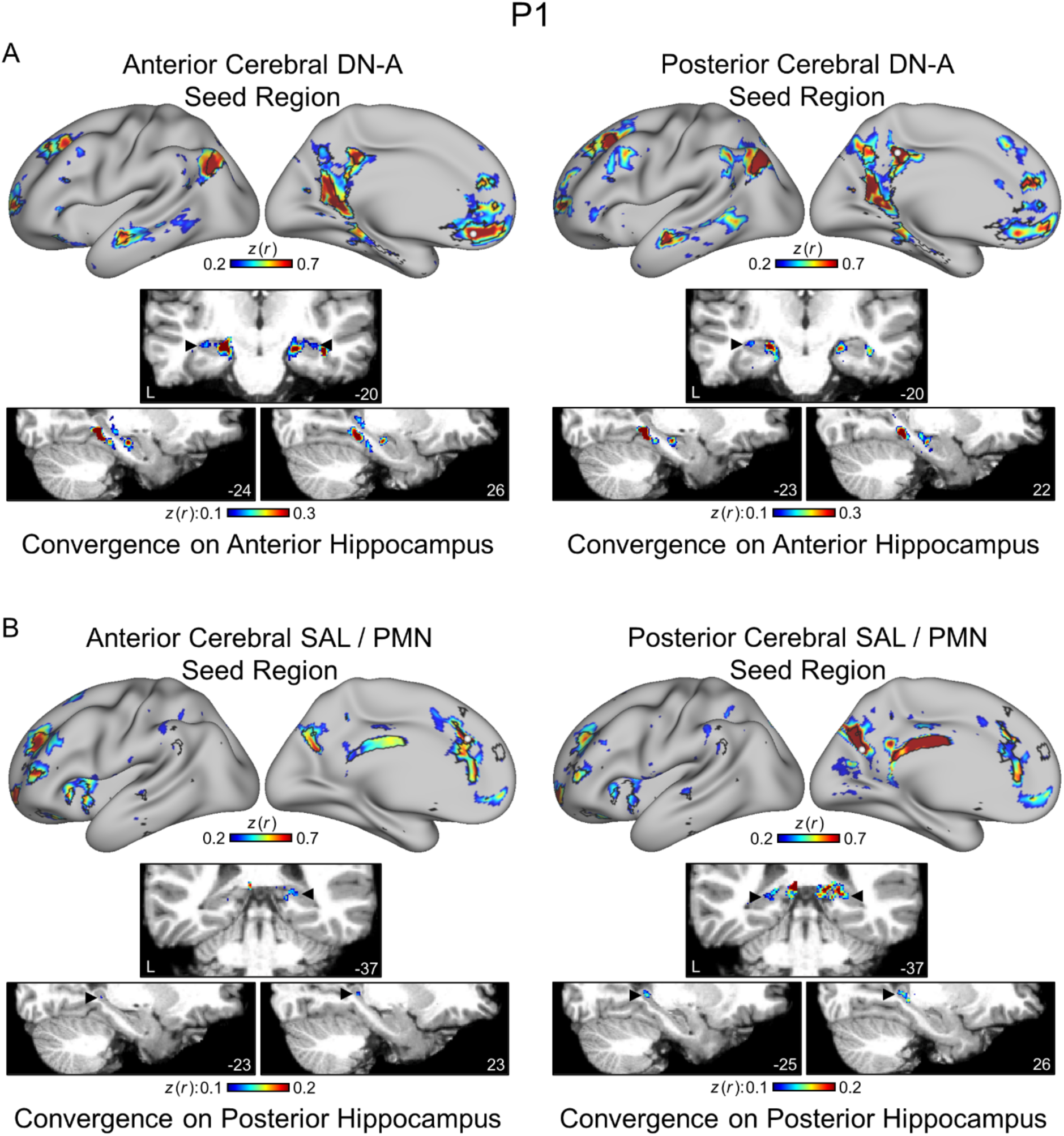
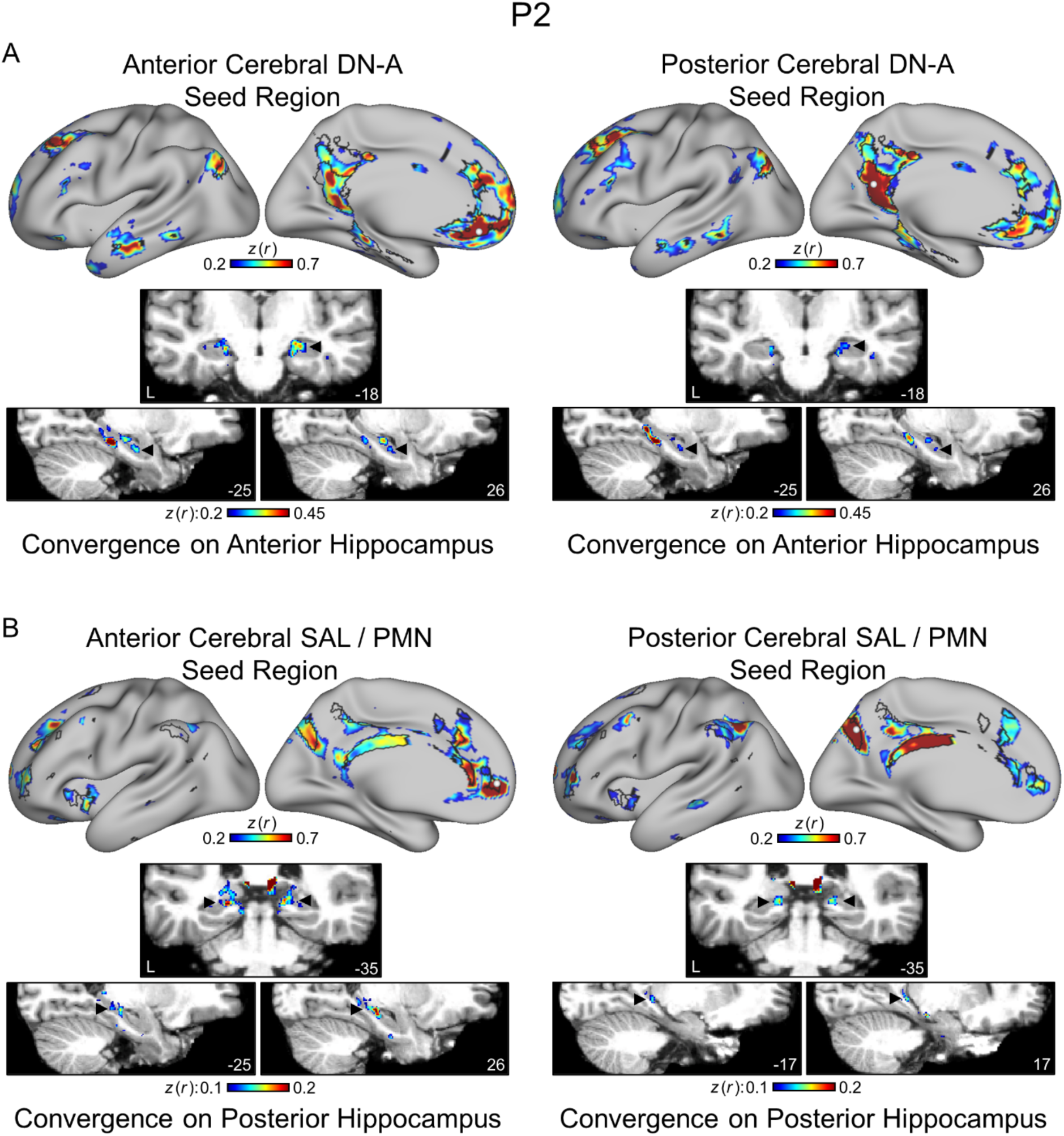
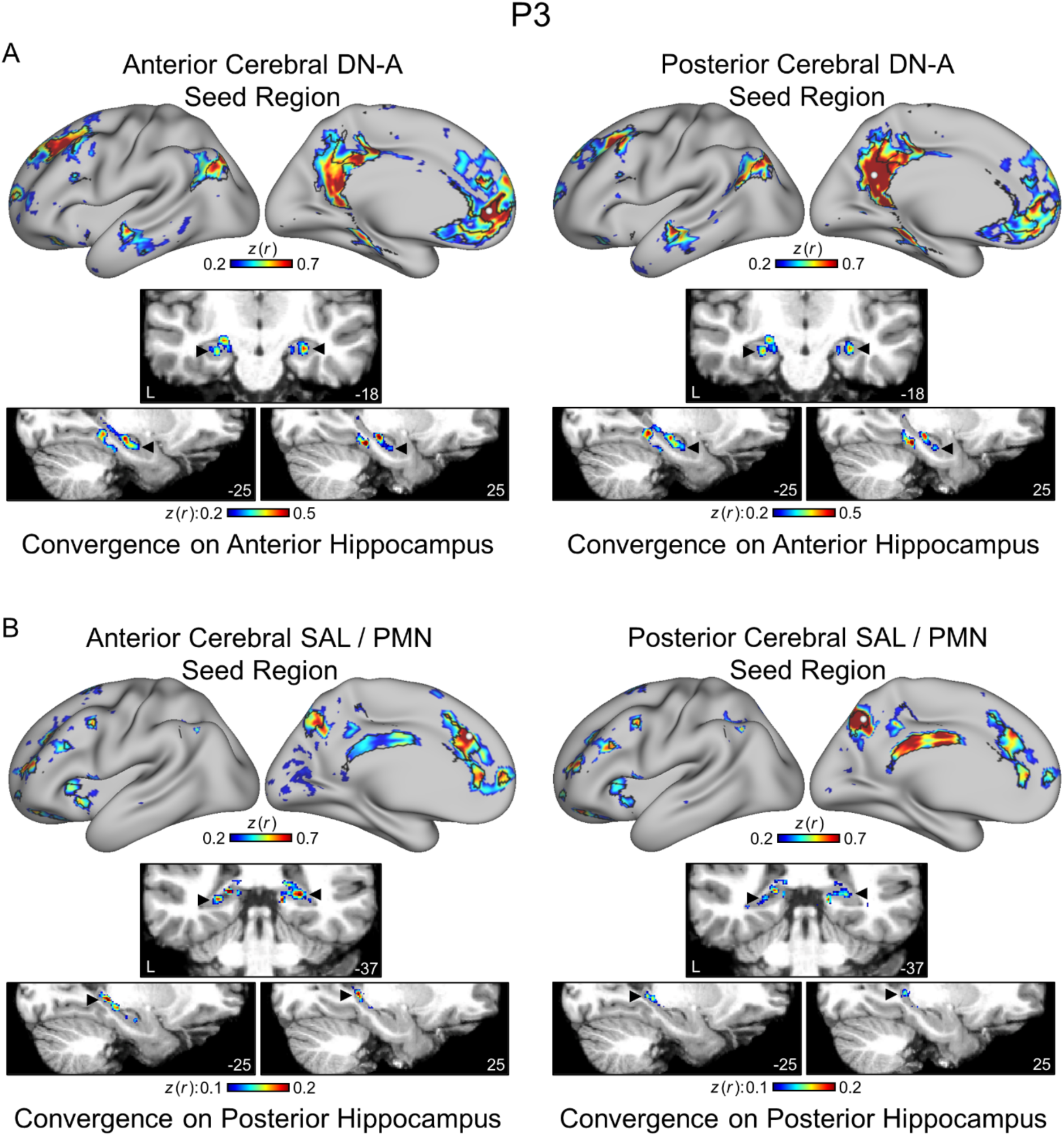
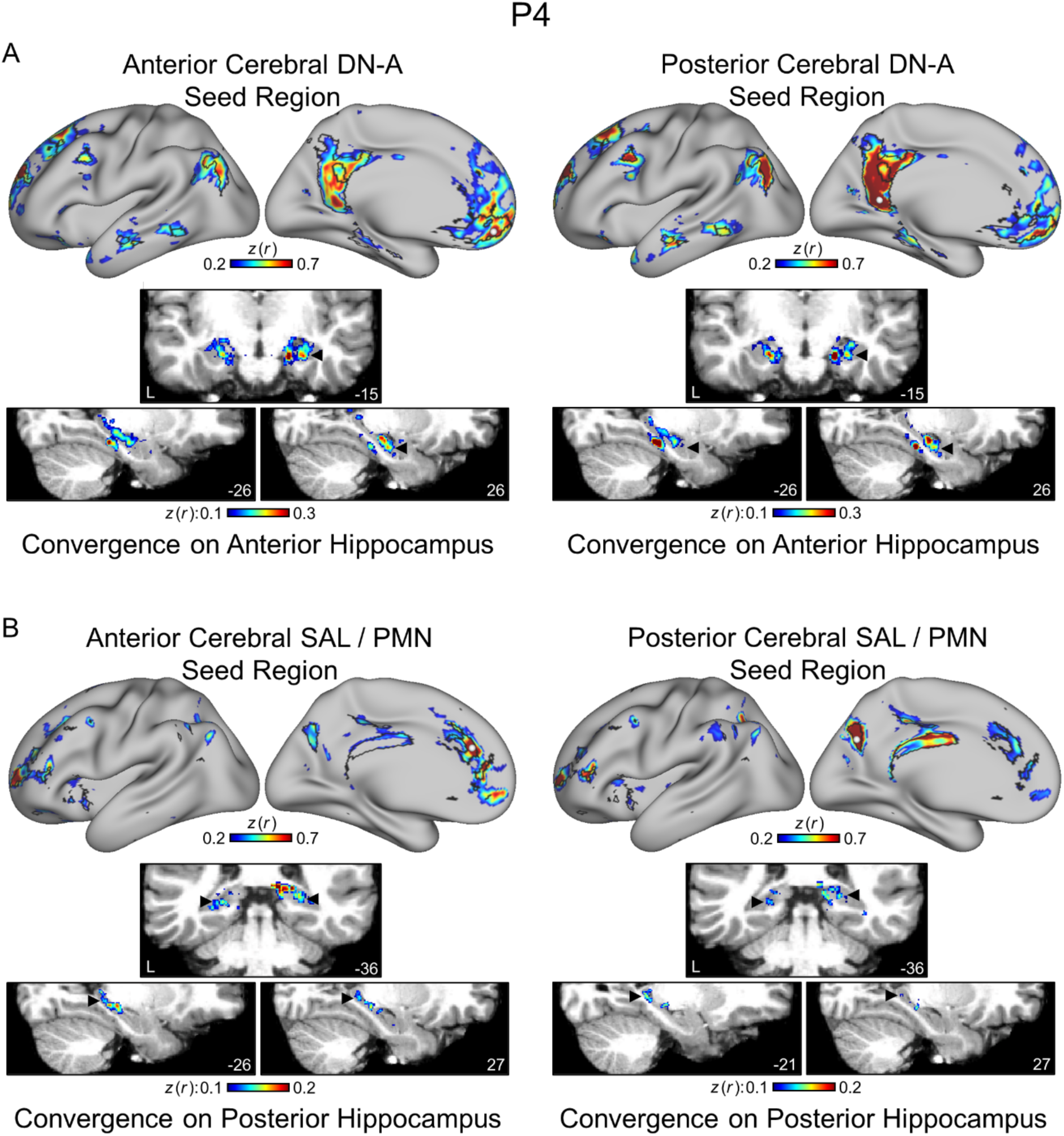
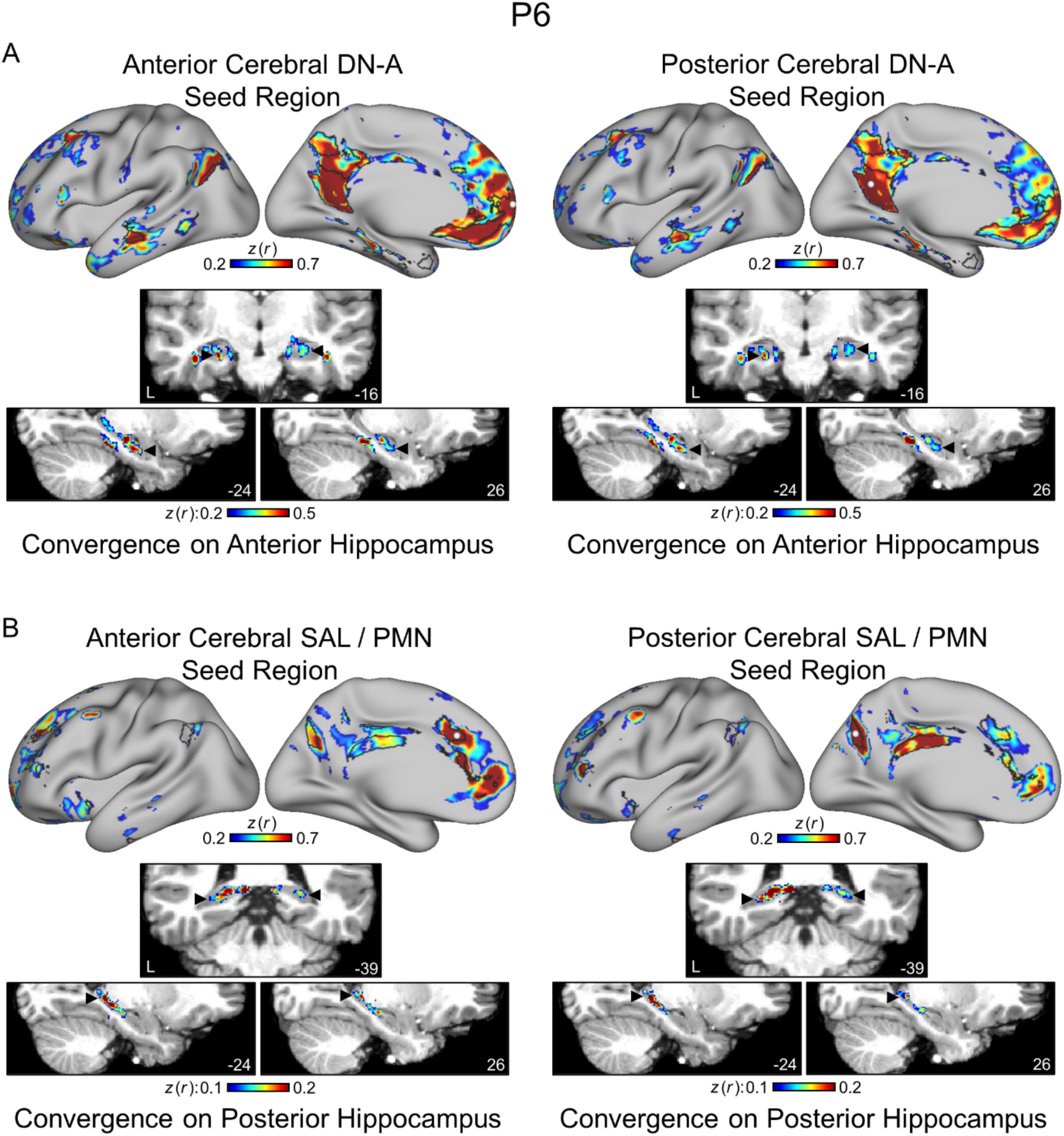
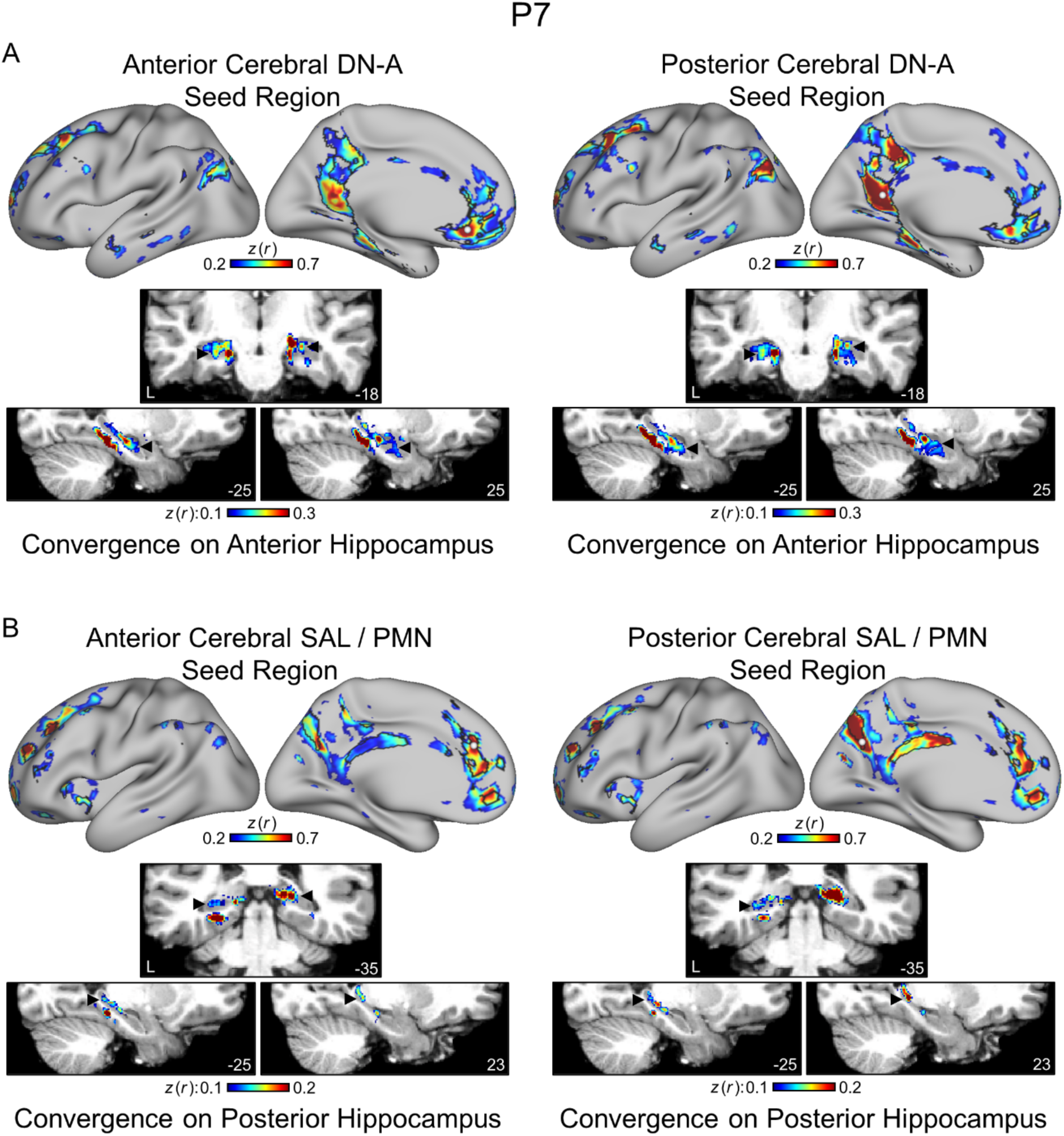
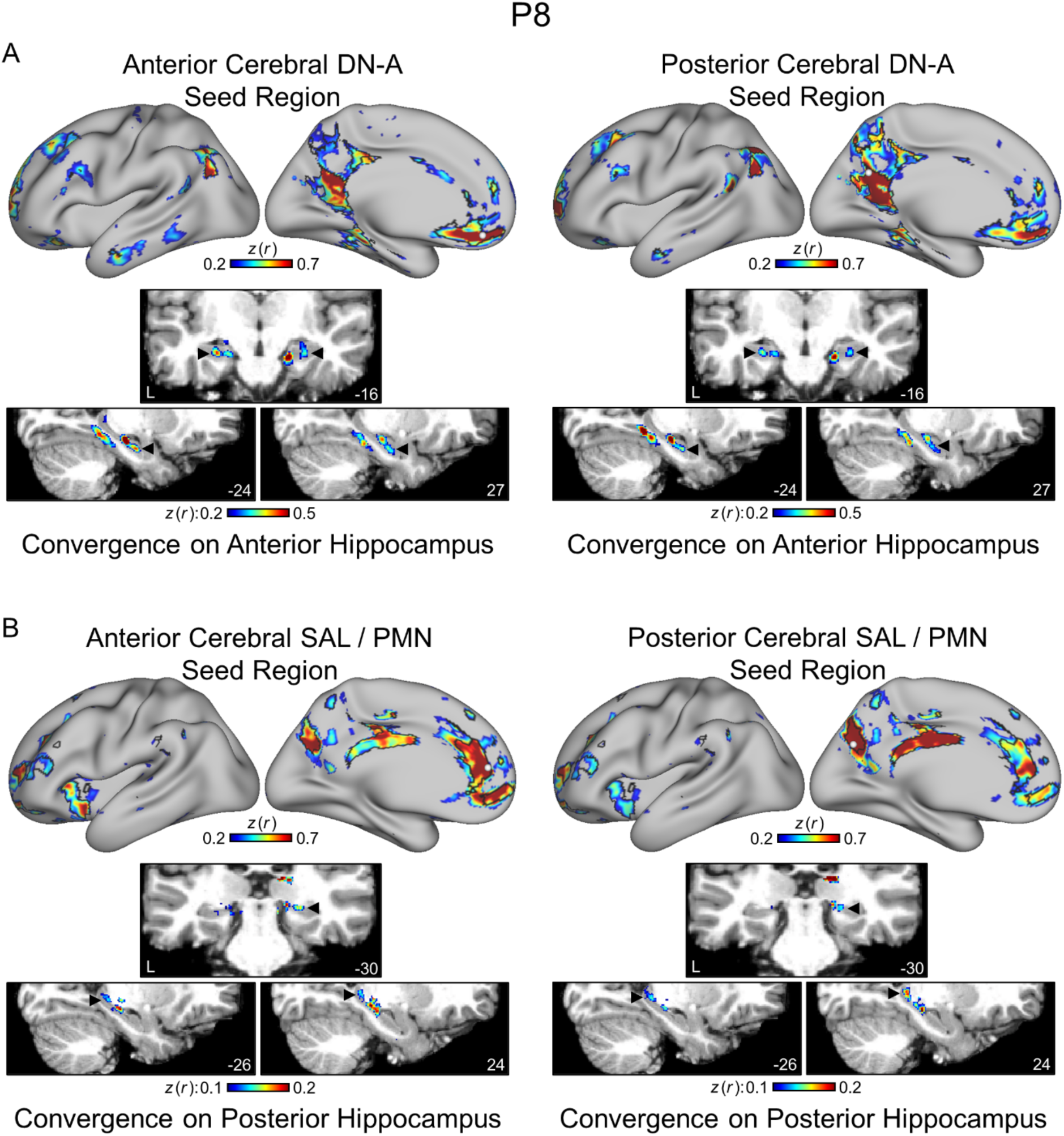
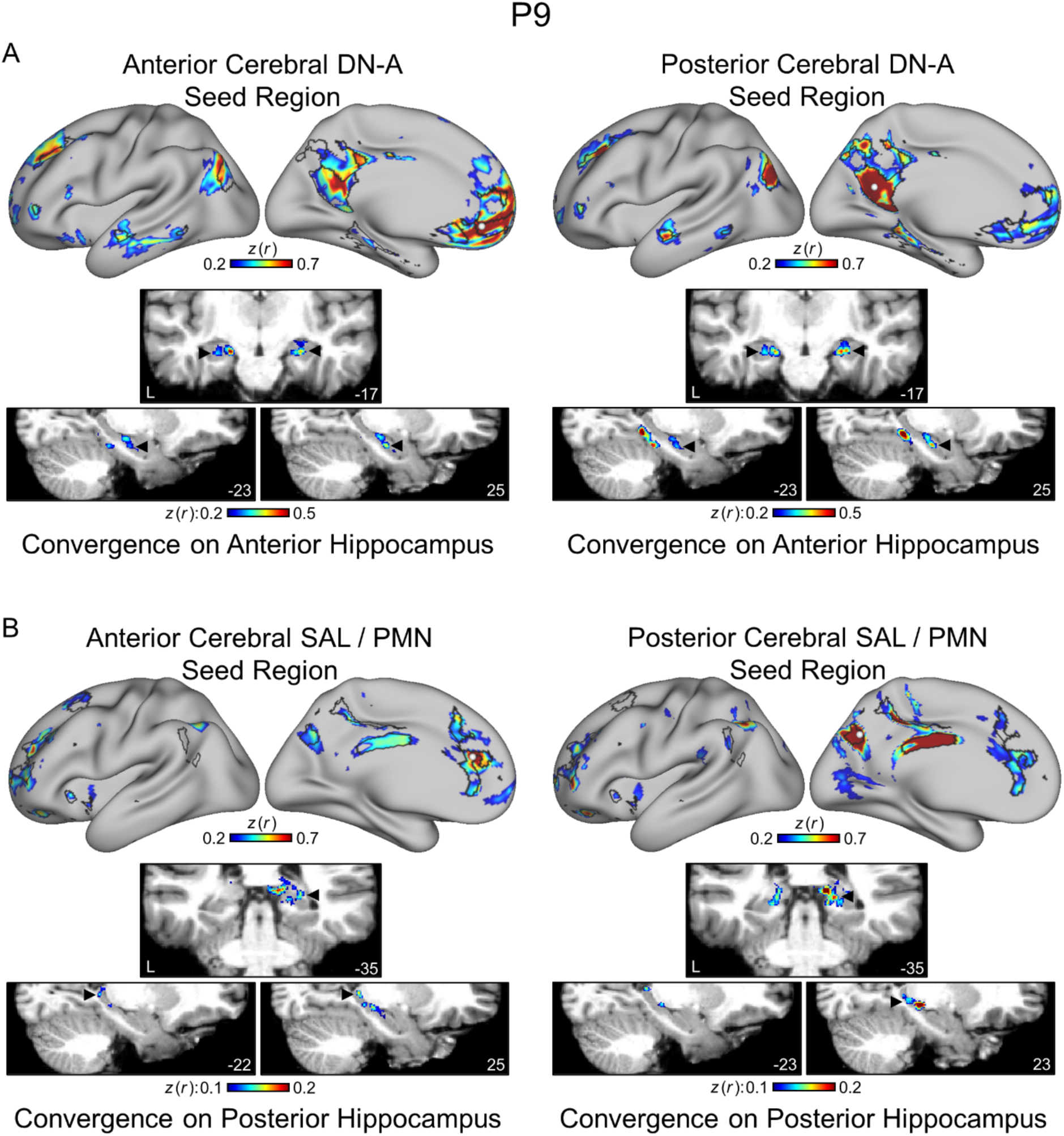
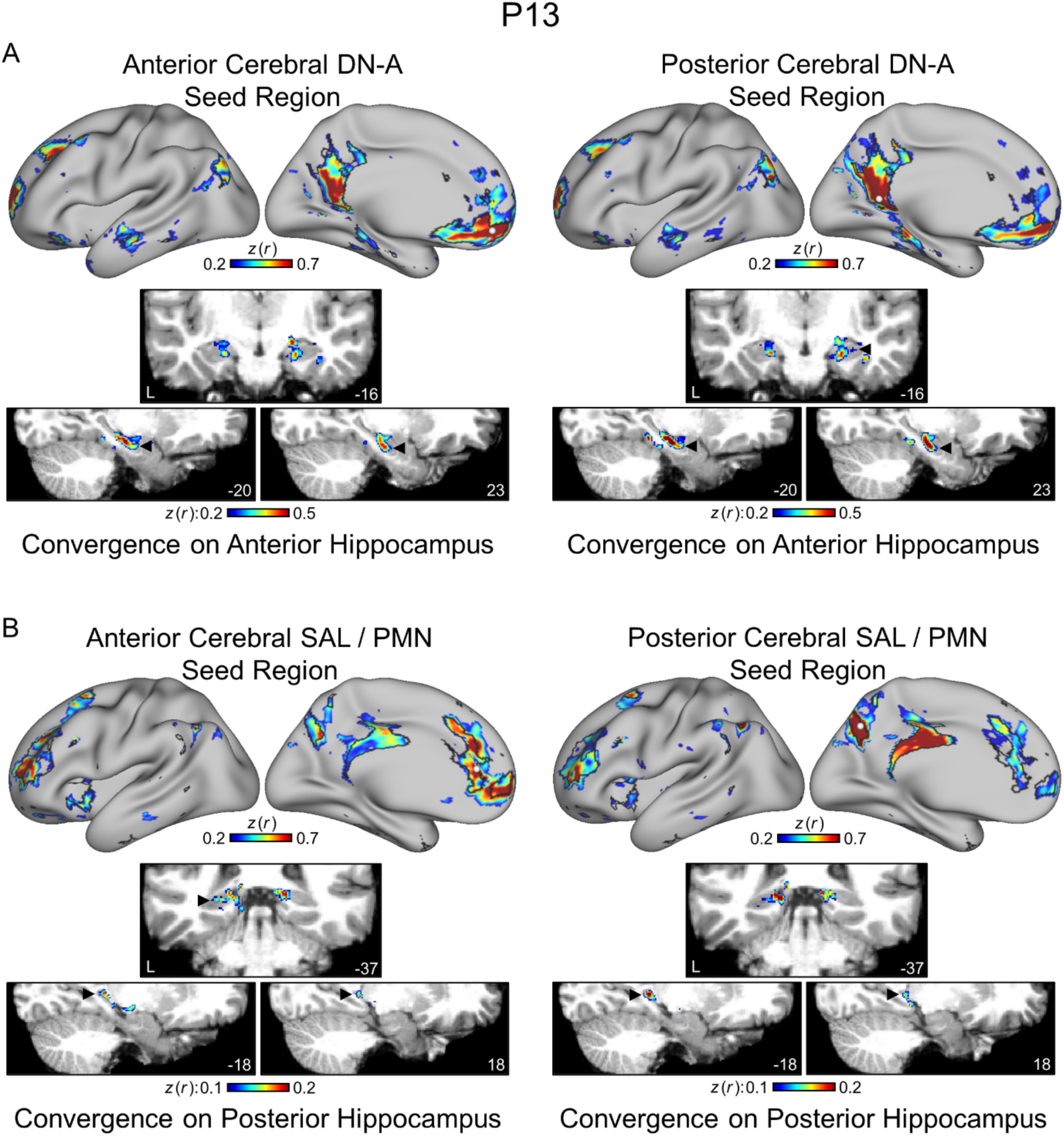
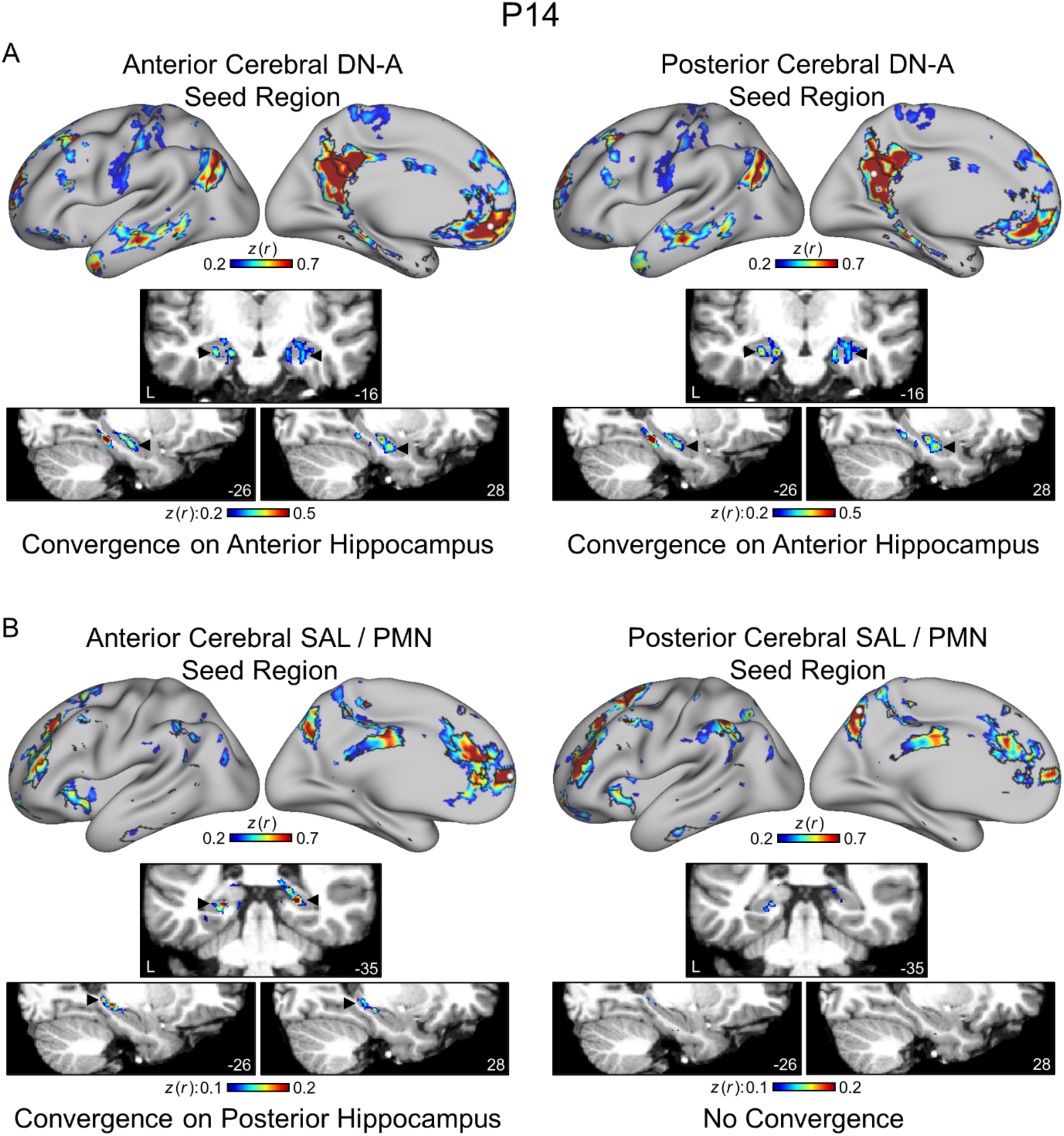
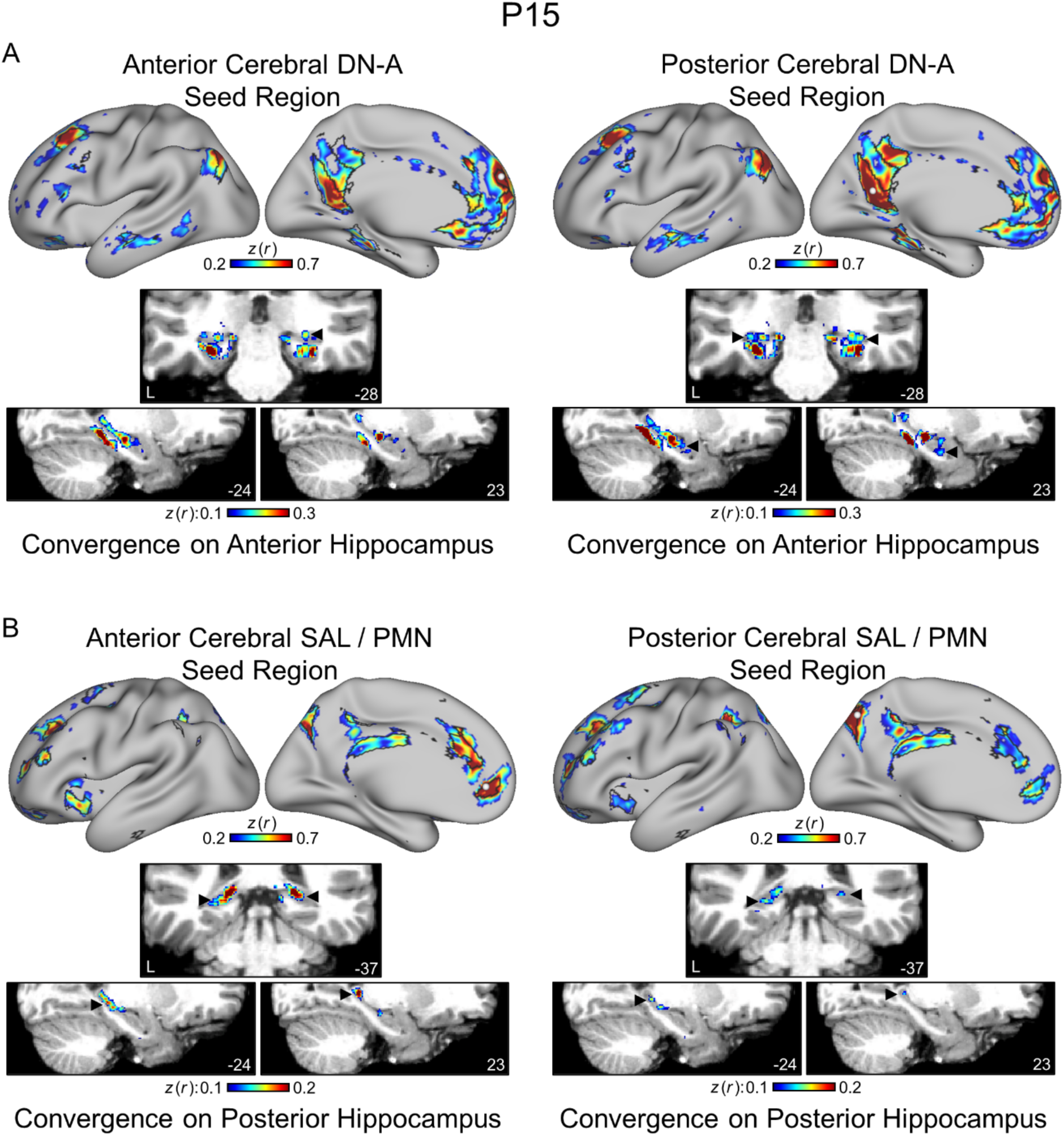
Unbiased seed-region based functional connectivity shows convergent cerebral network coupling to voxels within the hippocampus. Functional connectivity from a seed region on the cerebral surface to both the cerebral surface and the dilated hippocampal region in the volume. Each figure shows the maps for one participant. **(A)** Seed regions within the ventromedial prefrontal and retrosplenial portions of the DN-A network produced convergent patterns of functional connectivity in the anterior hippocampus (indicated by arrows). **(B)** Seed regions within the ventromedial prefrontal / anterior cingulate and posteromedial portions of the SAL / PMN network resulted in convergent functional connectivity within the posterior hippocampus (indicated by arrows).

**Supplemental Figure 16.**
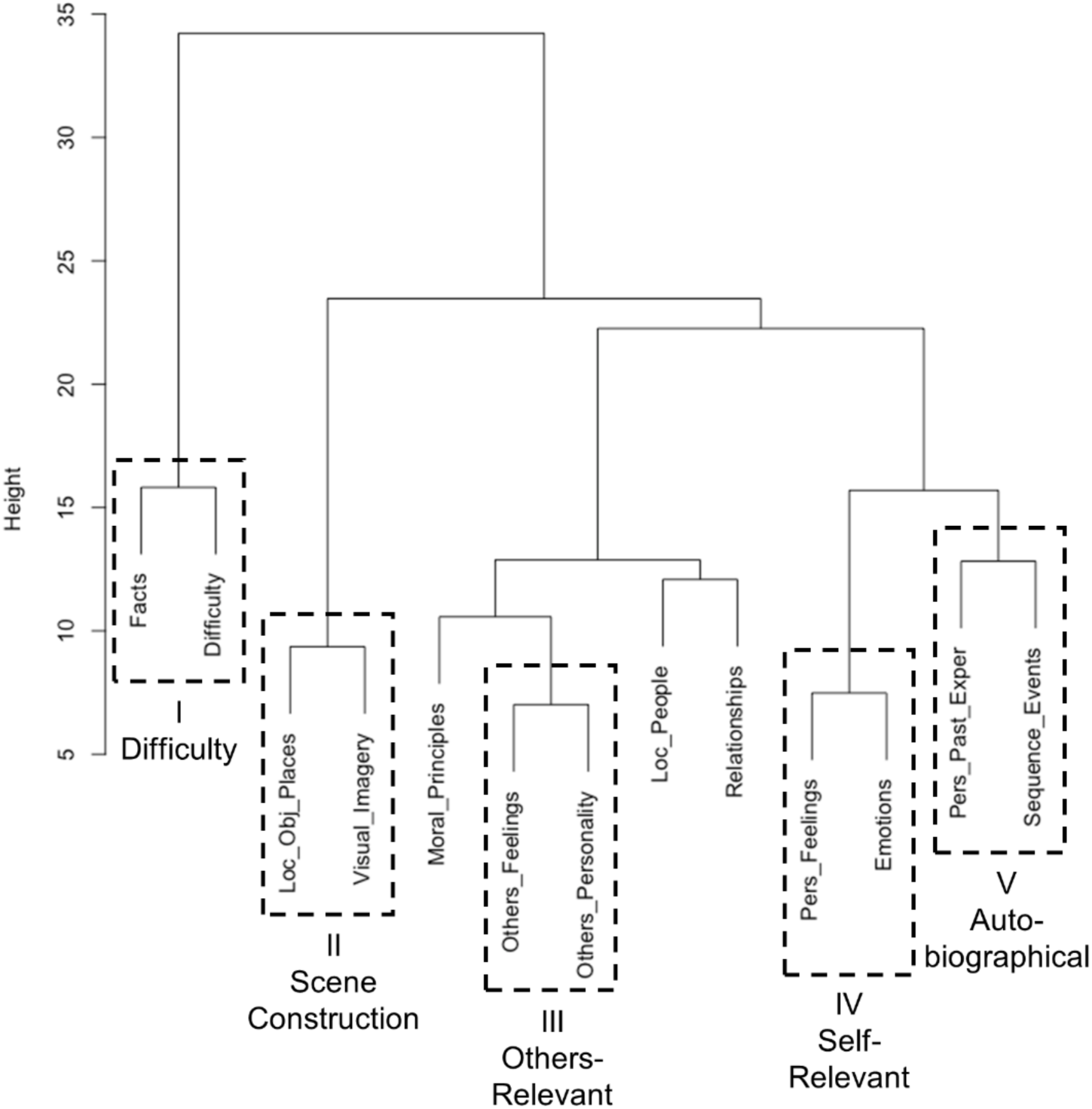
Episodic Projection trial-level strategy ratings cluster into five composite scores. Hierarchical clustering of Episodic Projection task trial-level strategy data using Ward’s minimum variance method. This clustering, and all subsequent analyses using these strategies, included only those 180 questions and 13 strategies shared with DiNicola et al. (2023a). All five of the clusters originally identified (and heuristically labeled I - difficulty, II - scene construction, III - others-relevant, VI - self-relevant, and V - autobiographical) were replicated in this new data set. The clusters were used as defined here for all analyses.

**Supplemental Figure 17.**
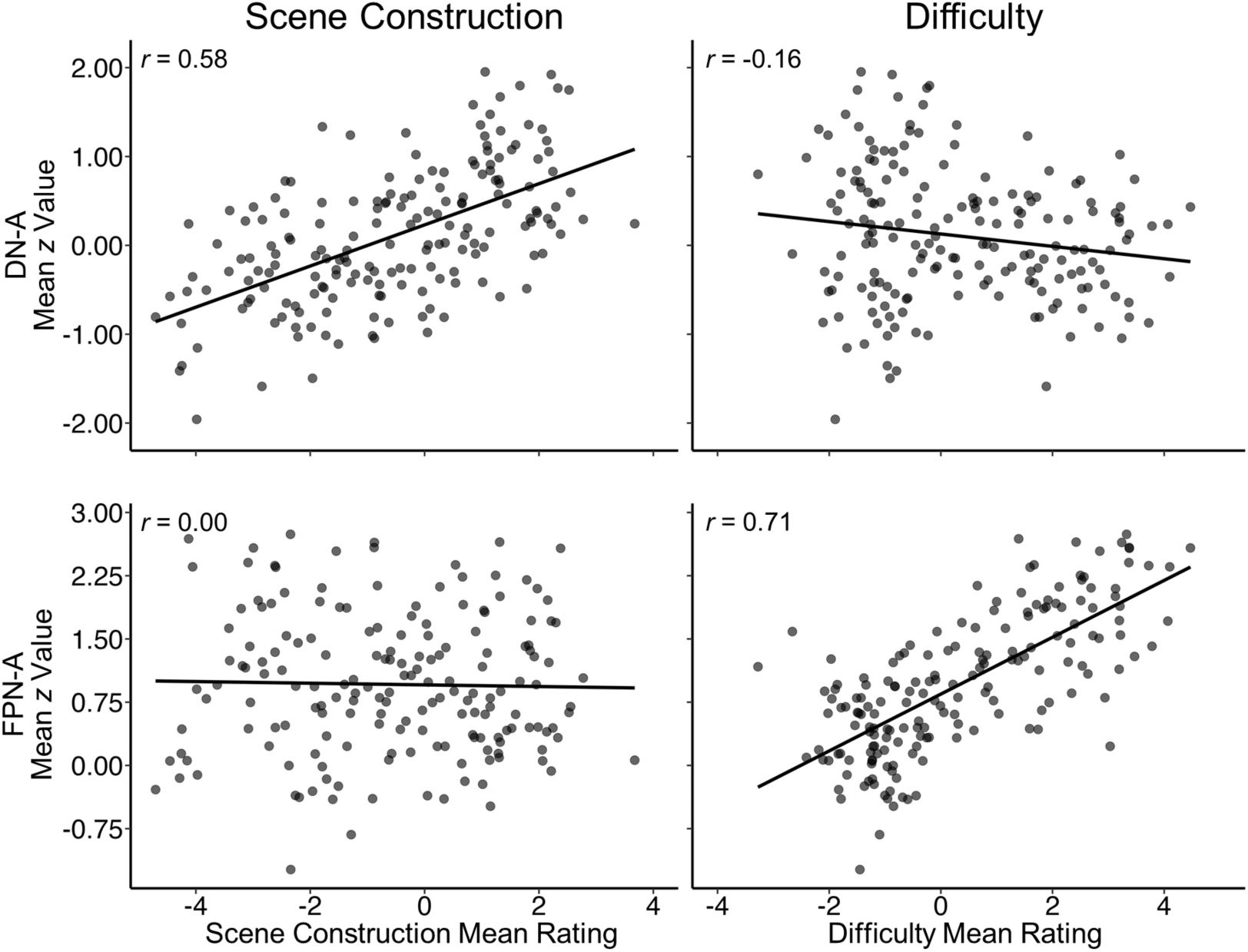
Cerebral networks DN-A and FPN-A differentially associate with behavioral estimates of scene construction and difficulty. Trial-level mean *z*-values within cerebral networks DN-A and FPN-A are correlated with behavioral estimates of scene construction and difficulty. Each point represents the mean activity value from all 11 MRI participants and each behavioral estimate reflects the mean composite score from 25 or more independent behavioral participants. DN-A is strongly correlated with scene construction but not difficulty, while FPN-A displays the opposite pattern of strong correlation with difficulty but not scene construction. This set of results prospectively replicates findings from DiNicola et al. (2023a) in independent neuroimaging and behavioral data. Note that in DiNicola et al. (2023a), FPN-A was labeled FPN-B. Here FPN-A is labelled to be consistent with work using the MS-HBM approach to network identification (e.g., DiNicola et al. 2023b, Du et al. 2023,.

**Supplemental Figure 18.**
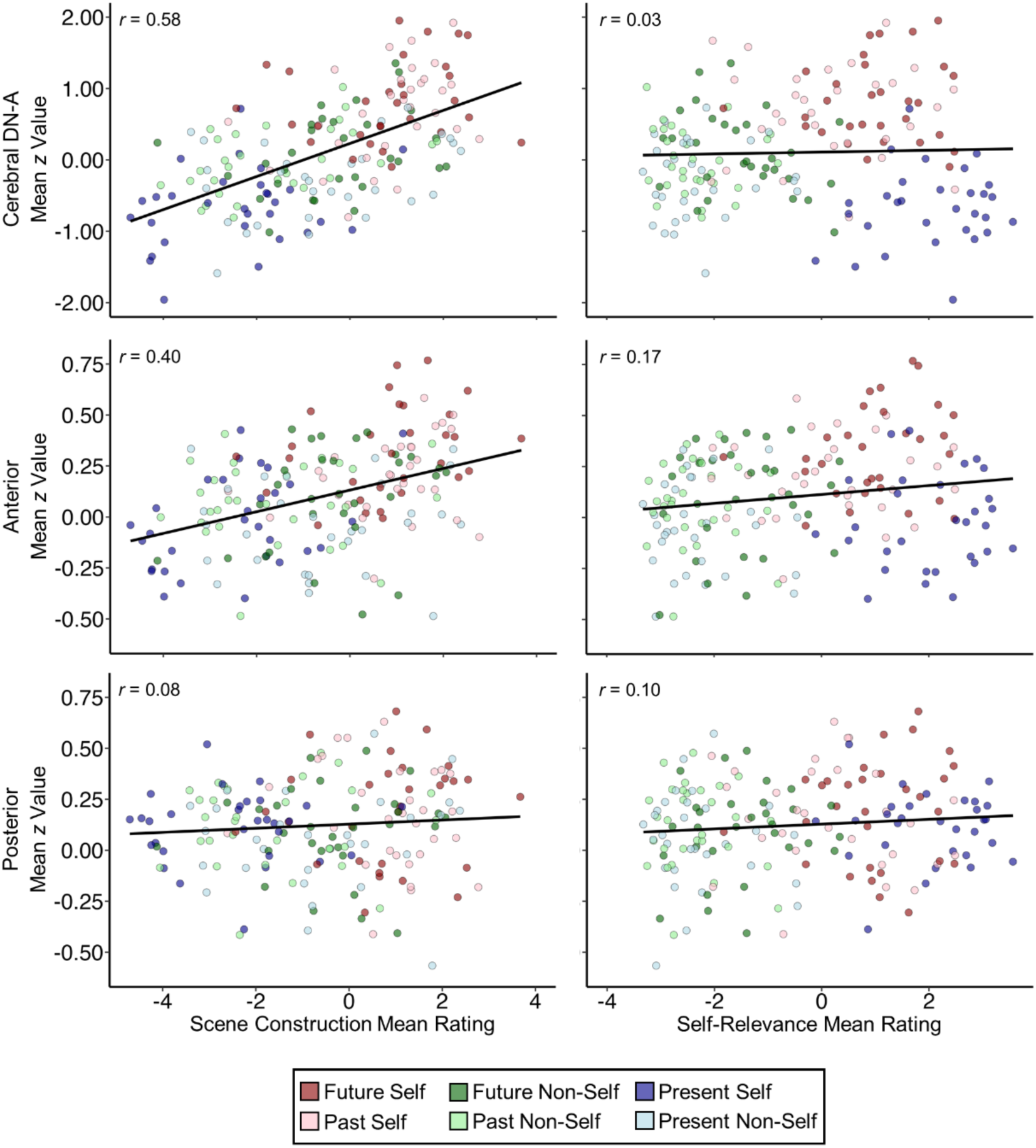
The anterior hippocampus tracks scene construction more than self-relevance during the Episodic Projection task. Trial-level mean *z*-values within cerebral network DN-A and the anterior and posterior hippocampal regions are correlated with the behavioral estimates of scene construction (left) and self-relevance (right). Each point represents the mean activity value from all 11 MRI participants and each behavioral estimate reflects the mean composite score from 25 or more independent behavioral participants. Each row reflects a different region (top, cerebral DN-A; middle, anterior hippocampus; bottom, posterior hippocampus). The points in the scatter plots are colored to show the original Episodic Projection task conditions (legend at bottom)(see DiNicola et al. 2023a for further description). Both DN-A and the anterior hippocampal region display strong, significant (*p* < 0.001) correlations with scene construction, while the posterior hippocampal region is not significantly correlated with scene construction (*p* = 0.27). Similarly, only anterior hippocampal region activity was significantly correlated with self-relevance (*p* < 0.05), and that correlation was weaker than for scene construction. Note that, even though both Future Self / Past Self and Present Self trials have similarly high self-relevance behavioral estimates, these trials show distinctly different activity levels in cerebral DN-A and the anterior hippocampal region. This is not the case for the relationship with scene construction, where across all conditions the trial-level activity levels are predicted well by behavioral estimates, and all fall along the same line, independent of condition. Even activity during control conditions (Future Non-Self, Past Non-Self, and Present Self, as in DiNicola et al. 2020) continues to track scene construction in cerebral DN-A and the anterior hippocampal region.

**Supplemental Figure 19.**
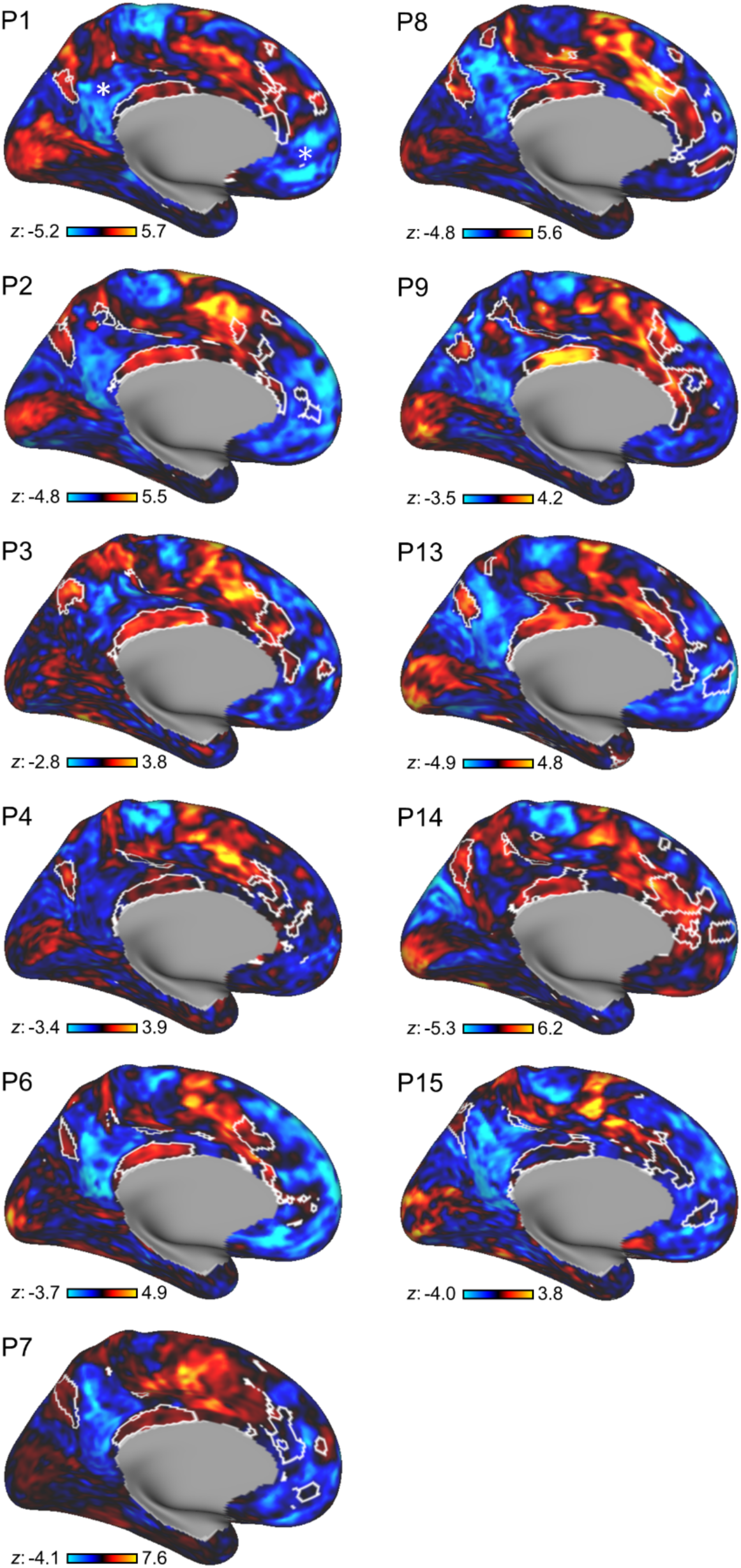
The visual oddball detection effect robustly dissociates the SAL / PMN from regions traditionally associated with the historically defined Default Network. Extending from Du et al. (2023), inflated surfaces display maps of the increases (red/yellow) and decreases (blue) in response for the visual oddball detection effect. No threshold is applied to allow full visualization of the effect in both directions. Images display all individual participants included in the present paper. The white outlines are the outline for the *a priori*-defined SAL / PMN network. Notice that the visual oddball detection effect increases response broadly across the SAL / PMN network and extends into adjacent regions (that are often part of the CG-OP network). The strong decreases span multiple networks including DN-A and DN-B (indicated by * on P1) that are canonical regions within the historically defined “Default Network.” This separation in the cerebral cortex echoes the dissociation observed in the hippocampus between the anterior and posterior hippocampal regions with the anterior hippocampus tracking the pattern observed for the cerebral DN-A network and the posterior hippocampus tracking the cerebral SAL / PMN network.

**Supplemental Figure 20.**
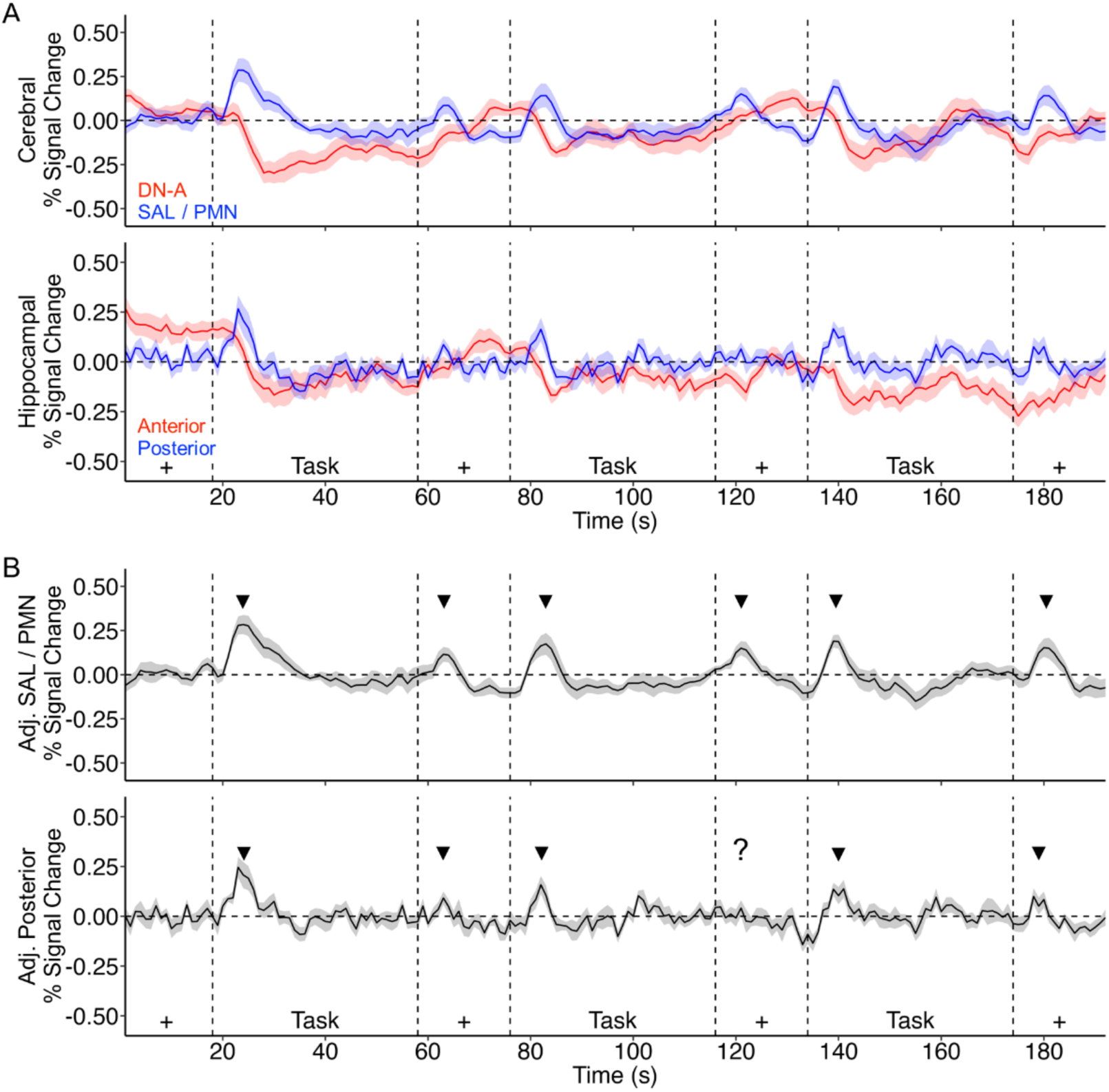
The cerebral SAL / PMN network and the posterior hippocampus transiently respond to oddball targets and task transitions. The present figure replots main text Fig. 4F with and without regression of the signal from cerebral DN-A. **(Top)** Time courses of the Blocked Visual-Motor task are shown for the cerebral SAL / PMN network and for the posterior hippocampal region both without signal regression of DN-A. The dashed lines indicate the transitions between blocks with the notation at the bottom indicating the block types (Task or fixation, +). The arrows indicate the transient responses at the block transitions. **(Bottom)** Adjusted time courses as shown in main text Fig. 4F of the Blocked Visual-Motor task are shown for the cerebral SAL / PMN network and for the posterior hippocampal region with DN-A signal regressed. The arrows again indicate the transient responses at the block transitions.

## Notes

### Competing Interest Statement

The authors have declared no competing interest.

